# Predicted distribution of a rare and understudied forest carnivore: Humboldt martens (*Martes caurina humboldtensis*)

**DOI:** 10.1101/2021.02.05.429381

**Authors:** Katie Moriarty, Joel Thompson, Matthew Delheimer, Brent Barry, Mark Linnell, Taal Levi, Keith Hamm, Desiree Early, Holly Gamblin, Micaela Szykman Gunther, Jordan Ellison, Janet S. Prevéy, Jennifer Hartman, Ray Davis

## Abstract

**Background:** A suite of mammalian species have experienced range contractions following European settlement and post-settlement development of the North American continent. For example, while North American martens (American marten, *Martes americana*; Pacific marten, *M. caurina*) generally have a broad range across northern latitudes, local populations have experienced substantial reductions in distribution and some extant populations are small and geographically isolated. The Humboldt marten (*M. c. humboldtensis*), a subspecies of Pacific marten that occurs in coastal Oregon and northern California, was recently designated as federally threatened in part due to its reduced distribution. To inform strategic conservation actions, we assessed Humboldt marten occurrence by compiling all known records from their range.

**Methods:** We compiled Humboldt marten locations since their rediscover to present (1,692 marten locations, 1996-2020). We spatially-thinned locations to 500-m to assess correlations with variables across contemporary Humboldt marten distribution (n=384). Using maximum entropy modeling (Maxent), we created distribution models with variables optimized for spatial scale; pre-selected scales were associated with marten ecology (50 to 1170 m radius). Marten locations were most correlated with abiotic factors (e.g., precipitation), which are unalterable and therefore uninformative within the context of restoration or management actions. Thus, we created variables to focus on hypothesized marten habitat relationships, including understory conditions such as predicted suitability of shrub species.

**Results:** Humboldt marten locations were positively associated with increased shrub cover (salal (*Gautheria shallon*), mast producing trees), increased pine (*Pinus sp*) overstory cover and precipitation at home-range spatial scales, areas with low and high amounts of canopy cover and slope, and cooler August temperatures. Unlike other recent literature on the species, we found little evidence that Humboldt marten locations were associated with old growth structural indices, perhaps because of a potential mismatch in the association between this index and shrub cover. As with any species distribution model, there were gaps in predicted distribution where Humboldt martens have been located during more recent surveys, for instance the southeastern portion of Oregon’s coast range. Conservation efforts and our assessment of potential risks to Humboldt marten populations would benefit from additional information on range extent, population sizes, and fine-scale habitat use. Like many rare and lesser-known species, this case study provides an example of how limited information can provide differing interpretations, emphasizing the need for study-level replication in ecology.

## Introduction

Modeling predicted distributions of rare or declining species can direct conservation efforts, but creating accurate predictions is challenging for a multitude of reasons. Contemporary location information may associate the species with conditions that were unaffected by prior agents of population decline, but not with favored characteristics where the species resided prior (Caughley 1994). For instance, bison (*Bison bison*) were historically widely distributed throughout western North America with herd sizes up to tens of millions of individuals (Shaw 1995), but species distribution models based on their contemporary distribution might suggest that they are associated with conditions present in Yellowstone National Park, such as extremely cold winters and thermal geysers. Instead, in this example, a logical conclusion is that bison in this area received less persecution. Nonetheless, a range-wide and broad-scale contemporary species distribution model ignorant of their biology and the causes of bison declines would misrepresent predicted habitat; creating more thermal geysers would not equate to more bison.

For populations that have declined or are currently declining, modeling species distributions may be complicated by additional factors – for example, contemporary locations may not include all areas where a species could physically colonize in the future, if the species is constrained by mobility or time (Barry et al. In review) or barriers that were not in place prior to the species’ decline (e.g., human development; Brown et al. 1996). Species may also require resources at multiple spatial scales, during different seasons, or for different durations of time – a female desert bighorn sheep (*Ovis canadensis nelsoni*) requires high nutritional-value vegetation during spring when providing for young (fine spatial extent, short duration), a water source within her range during the heat of the summer (large spatial extent, short duration), and immediate cover from predation year-round (fine spatial extent, long duration; e.g., Gedir et al. 2020; Hoglander et al. 2015). As such, limited observations in space or time may miss the conditions necessary for population persistence. Despite challenges for understudied species (Raphael & Molina 2007), spatial models can predict conditions of suitability that allow for species’ occurrence, which could identify areas of potential occupancy that are unknown due to lack of survey effort (Sofaer et al. 2019).

Modeling habitat and predicted distribution of North American marten populations (American marten, *Martes americana*; Pacific marten, *M. caurina*) is particularly challenging, due to presumed selection of both broad-scale landscape features (1st order selection sensu Johnson 1980) and fine-scale features within home ranges (4th order selection; e.g., Minta et al. 1999). Martens are small-bodied (600-1300g) carnivores that occupy large home ranges relative to their body size and circumnavigate the perimeter of their territory often (e.g., every 4-5 days, Balharry 1993; Moriarty et al. 2017). To fulfill some life history requirements, martens select fine-scale forest elements that facilitate foraging, daily resting and seasonal denning activities, and allow them to avoid predators (Andruskiw et al. 2008; Cheveau et al. 2013; Joyce 2013; Shirk et al. 2012). Simultaneously, fulfillment of other life history requirements (e.g., size and positioning of home ranges relative to intraspecific competitors or potential mates) requires selection for broader-scale forest attributes such as cohesive and complex vegetation cover (Hargis et al. 1999; Potvin et al. 2000); as such, martens have been highlighted as a species benefiting from multi-scale habitat selection models (Bissonette & Broekhuizen 1995; Shirk et al. 2014; Wasserman et al. 2010).

Humboldt martens (*M. c. humboldtensis*) are a subspecies of Pacific marten that historically occurred throughout coastal forests of northern California and Oregon. Contemporary range-wide surveys suggest that distribution of Humboldt martens is substantially reduced compared to historical representations (Moriarty et al. 2019; Zielinski et al. 2001). Consequently, Humboldt martens were listed as Endangered under the state of California’s Endangered Species Act (CDFW 2019) and as threatened under the federal Endangered Species Act as a “coastal distinct population segment of Pacific martens” (USFWS 2020). Evaluating current and future Humboldt marten distribution is urgently needed for conservation planning, yet modeling their distribution is constrained by a paucity of ecological information (e.g., population sizes and extents) and apparent non-stationary associations with vegetation among populations. For example, Humboldt martens have been associated with a diversity of forest types across their distribution, including mature forests with high canopy cover, mature forests with dense shrub understories (Slauson et al. 2007), young forests (<80 years old) with modest canopy cover and relatively small diameter trees with dense shrub cover (Eriksson et al. 2019; Moriarty et al. 2019), and forests in serpentine soils with sparse canopy cover and dense shrub cover. A dense and spatially-extensive shrub layer appears to be an attribute shared by each of these studies (Eriksson et al. 2019; Gamblin 2019; Moriarty et al. 2019; Slauson et al. 2007).

A previous range-wide Humboldt marten distribution model Slauson et al. (2019b) emphasized a strong correlation between Humboldt marten occurrence and an “old-growth structural index” (OGSI) variable, which is a composite index of factors considered common to old-growth forests in the region, including density of large live trees, snags and downed wood, and diversity of tree sizes (Ohmann 2012). However, the model relied on modest number of detections from 1996–2010 with poor coverage outside a portion of northern California (USFWS 2019). Since 2010, we initiated large-scale surveys for Humboldt martens that greatly increased the spatial extent and number of Humboldt marten detections in both California and Oregon (e.g., Barry 2018; Gamblin 2019; Linnell et al. 2018; Moriarty et al. 2019). Further, recent research efforts appear to contradict the finding that Humboldt martens are strongly associated with old-growth forests, particularly in Oregon (e.g., Eriksson et al. 2019), suggesting a potential mismatch in previously-predicted associations between OGSI and Humboldt marten distribution. Such a mismatch could lead to a “wicked problem” by focusing management or restoration in areas that would not benefit the species (Gutiérrez 2020). For example, a landscape connectivity model for Humboldt martens determined both “habitat cores” and connectivity between cores based almost exclusively on OGSI values (Schrott & Shinn 2020).

Here, our objective was to create a contemporary range-wide model of predicted Humboldt marten distribution facilitated by including recent location data collected from broad-scale randomized surveys throughout the historic range, combined with more recent and accurate vegetation layers (e.g., shrub layers). Our goal was to predict factors contributing to probabilities of Humboldt marten spatial use and to highlight areas for future surveys and conservation efforts.

## Materials & Methods

### Study Area

Humboldt martens are thought to occur in four populations in California and Oregon, represented as the Central Coastal Oregon, Southern Coastal Oregon, California-Oregon Border, and Northern Coastal California populations (USFWS 2019; Fig. 1). Data were collected in Oregon in both near-coastal and montane areas (Oregon Coast Range), where dominant forest types include Sitka spruce (*Picea sitchensis*) on the coast and western hemlock (*Tsuga heterophylla*) (Franklin & Dyrness 1973). Sitka spruce zones were mild (annual average temperature 10-11 °C), averaging 200-300 cm of annual precipitation interspersed with regular fog and cloud cover. Western hemlock zones, often co-dominated by Douglas-fir (*Pseudotsuga menziesii*), were are also wet (150-300 cm annual precipitation) and somewhat cooler (7-10 °C) environments but experience drier summers (6-9% of total precipitation) with fairly extensive summer fog and low cloud cover (Dye et al. 2020). Surveys in California similarly occurred in both coastal and montane areas (Klamath Mountains, California Coast Range), with dominant forest types including redwood (*Sequoia sempervirens*) along the coast and Douglas-fir or mixed-conifer in the mountains (Whittaker 1960). These areas included a mix of coniferous and hardwood forest with a spatially-extensive shrub understory and often received substantial precipitation (100-300 cm annual precipitation) with cooler (7-10 °C) environments but with drier summers dominated with fog and low cloud moisture (Rastogi et al. 2016). Dominant forest types include tanoak (*Notholithocarpus densiflora*) in the southeastern portion of the northern California study area. Common conifer species included western hemlock, Port Orford cedar (*Chamaecyparis lawsoniana*), and western redcedar (*Thuja plicata*). Common hardwood species included chinquapin (*Chrysolepis chrysophylla*), red alder (*Alnus rubra*), bigleaf maple (*Acer macrophyllum*), and Pacific madrone (*Arbutus menziesii*). Dominant shrub species throughout the range includes salal *(Gautheria shallon*), evergreen huckleberry *(Vaccinium ovatum*), Pacific rhododendron *(Rhododendron macrophyllum*), and red huckleberry *(V. parvifolium*).

**Figure 1.**
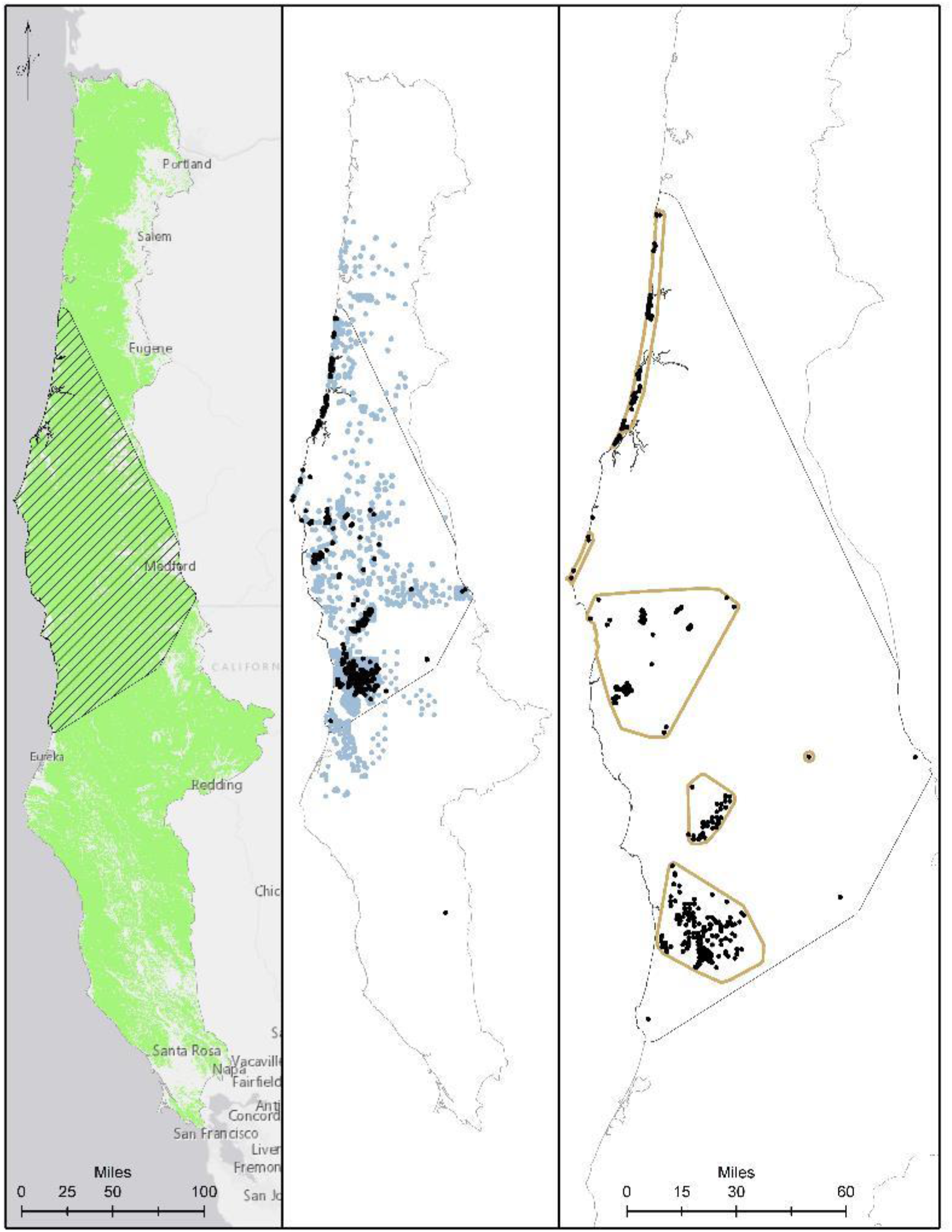
Our study area and modelling region for Humboldt martens (*Martes caurina humboldtensis*) included all of coastal Oregon and northern California. We modeled Humboldt marten predicted distributions in forested lands (Panel A, green mask) in 2 ecoregions [left]. We created a minimum convex polygon of known locations buffered by 10-km (hatched area). We compiled 1,692 known marten locations (icon color) from 5,143 surveyed sites with non-detections in light gray, collected during 1996-2020 (panel B). We spatially thinned locations to approximately 500m apart, prioritizing den and rest locations and resulting in 384 locations (black dots, panel C).

### Marten locations

We incorporated verifiable, contemporary, spatially-referenced Humboldt marten (hereafter, marten) locations collected during 1996-2020 into our model. We removed marten locations from our model sample that occurred in areas that disturbance (fire or harvest) between the data of observation and the date represented by our vegetation layers (2016). If there were multiple locations within the 500-m cell, we spatially-thinned locations prioritizing those in the order of: (1) rest and den locations from telemetry studies in the Central Coastal Oregon population (Linnell et al. 2018) and the Northern Coastal California population (Delheimer et al. In press; PSW 2019); (2) locations from scat dog detection surveys (Rogue Detection Teams, Rice, WA; Conservation Canines, University of Washington, Seattle, WA) from both Oregon populations (detailed methods within Moriarty et al. 2018; Moriarty et al. 2019); and (3) locations from baited camera and/or track plate surveys from all four populations in Oregon (Barry 2018; Moriarty et al. 2019) and California (Gamblin 2019; Slauson et al. 2012). We thinned locations to one within a 500-m cell, attempting to maximize distances between locations and induce spatial independence for modeling (Kramer-Schadt et al. 2013). We used presence-only data to incorporate telemetry-based information and detection dog surveys in areas that outside the geographic extent of baited and lured detection/non-detection surveys and because data on survey locations without detections and detection histories were not available for older surveys (e.g., <2014).

For the data in which the authors were responsible, our protocols were reviewed by the USDA Forest Service Research and Development or Humboldt State University’s Institutional for Use and Care Committee. We received approval for marten captures and treatment in the Central Coast (USDA FS R&D 2015-002). We received a waiver for camera trapping in Oregon by the same committee (USDA FS R&D 2017-005) and camera surveys in the Border population were under Humboldt State University 16/17.W.05-A. We obtained Scientific Take Permits for hair snares and samples collected through the Oregon Department of Fish and Wildlife (ODFW 119-15, 128-16, 033-16, 109-19, 107-20). Other verified survey data were provided by the US Fish and Wildlife Service with no additional information.

### Modeling approach

Our overall modeling approach incorporated our collated Humboldt marten locations, biotic and abiotic predictor variables, and randomly-generated background points (*n* = 10,000). We used a minimum convex polygon around Humboldt marten locations buffered by 10 km for the model training and testing (Fig. 1b). We chose a 10 km buffer because this was approximately the upper quartile of daily marten movement (Moriarty et al. 2017). We projected our model to available vegetation data from Gradient Nearest Neighbor (GNN) 2016 data supplied by the Landscape Ecology, Modeling, Mapping and Analysis lab (Bell et al. 2021; Bell et al. 2020), which included the coastal and Klamath level-3 eco-provinces (U.S. Environmental Protection Agency 2013). We applied a mask to only include areas typed for forested vegetation, which removed urban areas and water, similar to Davis et al. (2016). We summarized the range, average, and standard deviation for each variable at used marten locations, within the minimum convex polygon (MCP) boundary within a 10-km buffer around known marten locations, and the overall study area (Table 1, Fig. 1).

**Table 1.** Data ranges, means, and standard deviations for the model region, the contemporary Humboldt marten distribution, and at Humboldt marten locations. We depict individual layer statistics within our Humboldt marten (*Martes caurina humboldtensis*) model region in coastal Oregon and northern California. We display the variable, optimized spatial scale with a radius in meters, value range from the coastal ecoregions, means and standard deviation (SD) for the model region, minimum convex polygon around all known marten locations (MCP), and values from spatially thinned marten locations (n = 384), our layer source, and a description of that variable. We only considered variables with < 60% correlation in our final model (Table S2).

| Variable | Scale | Value Range | Model Region (Mean ± SD) | Minimum convex polygon (Mean ± SD) | Marten Locations (Mean ± SD) | Source | Description |
| --- | --- | --- | --- | --- | --- | --- | --- |
| Forest age, years | 270 | 0 – 712 | 95.5 ± 43 | 104.3 ± 49.4 | 109.8 ± 69.6 | 2016 GNN | Basal area weighted stand age based on field recorded or modeled ages of dominant/codominant trees |
| Canopy cover (%) | 1170 | 2 – 99 | 65.9 ± 13 | 66.4 ± 14 | 71.3 ± 18.6 | 2016 GNN | Canopy cover percentage of all live trees |
| Coastal proximity | 50 | 2 – 700 | 511.7 ± 193.1 | 516.3 ± 203.1 | 361.8 ± 197.9 | PRISM | Optimal path length from the coastline accounting for terrain blockage (Daly et al. 2008) |
| Diameter diversity index | 1170 | 26 – 811 | 433.9 ± 103 | 437.6 ± 111.7 | 459.4 ± 123.6 | 2016 GNN | Diameter diversity index - measure of stand structure based on tree densities in diff. DBH classes (x100) |
| Percent<br>downed wood | 270 | 0 – 797 | 69.3 ± 54.7 | 70.9 ± 50 | 68.5 ± 60.1 | 2016<br>GNN<br>(created) | Created within GNN to estimated percentage of large downed wood, a component of OGSi |
| Salal | 1170 | 0 – 100 | 35.7 ± 30.9 | 50.7 ± 32.3 | 72.7 ± 17.8 | Prevéy | Probability of *Gautheria shallon* species occurrence (Prevéy et al. 2020) |
| Masting<br>vegetation | 1170 | 0 – 72 | 5.9 ± 7.4 | 5.2 ± 6.7 | 9.3 ± 9 | 2016<br>GNN | Percent of stand basal comprised of tanoak (*Notholithocarpus densiflorus*; LIDE), giant chinquapin (*Castanopsis chrysophylla*; CHCH), coastal live oak (*Quercus agrifolia*; QUAG), canyon live oak (*Quercus chrysolepis*; QUCH), and California bay (*Umbellularia californica*; UMCA) (mast producing evergreen hardwoods, indicator of prey abundance) |
| Old growth<br>structural index | 50 | 0 – 100 | 32.7 ± 15.8 | 33.2 ± 16.1 | 33.8 ± 16.9 | 2016<br>GNN | Old-growth structure index based on abundance of large live trees, snags, down wood, and ddi |
| Percent pine | 1170 | 0 – 94 | 1.2 ± 3.5 | 1.5 ± 4.5 | 10.9 ± 20.1 | 2016<br>GNN | Percent of pixel basal area comprised of shore pine (*Pinus contorta*; PICO), Jefferey pine (*Pinus jeffreyi*; PIJE) and knobcone pine (*Pinus attenuata*; PIAT). We use this as an indicator of serpentine and coastal dune environments. |
| Percent slope | 1170 | 0 – 74 | 33.8 ± 10.9 | 36.2 ± 10.6 | 31.7 ± 15.8 | USGS<br>DEM | Percent slope in degrees |
| Precipitation | 1170 | 13 – 198 | $66.9 \pm 27$ | $70 \pm 30.1$ | $102.4 \pm 30.5$ | 2016 GNN | Average annual precipitation 1981-2010 (inches) |
| Large snag density | 742 | 0 – 48 | $4.9 \pm 4.3$ | $5.8 \pm 4.6$ | $6.9 \pm 4.9$ | 2016 GNN (created) | Created within GNN to estimated density of large snags, a component of OGSi |
| Temperature (August max) | 1170 | 8 – 24 | $16.5 \pm 2.3$ | $16.1 \pm 1.7$ | $16.4 \pm 1.7$ | PRISM | Average annual maximum temperature 1981-2010 (Celcius). |
| Topographic position index | 270 | -149 – 174 | $0.7 \pm 26.7$ | $1.1 \pm 28.8$ | $-0.3 \pm 28.6$ | USGS DEM | Topographic position index - difference of cell elevation with mean of all cells w/in 450 m radius |
| Large tree density | 1170 | 0 – 47 | $3.2 \pm 3.5$ | $4.4 \pm 4.2$ | $5.2 \pm 5.9$ | 2016 GNN (created) | Created within GNN to estimated density of large trees, a component of OGSi |
| Huckleberry | 1170 | 2 – 99 | $32.7 \pm 24.6$ | $39.1 \pm 26$ | $42.7 \pm 27.2$ | Prevéy | Probability of species occurrence for *Vaccinium ovatum* (created) |

### Biotic variables

We incorporated biotic variables into our models depicting forest structure and forest composition (e.g., tree and shrub species). We acquired variables representing forest structure through the 2016 version of GNN (Ohmann & Gregory 2002) and included forest age, canopy cover, and OGSI. We depicted forest age as the basal area-weighted age based on field-recorded or modeled ages of dominant and codominant trees. Canopy cover percent was calculated using methods in the Forest Vegetation Simulator (Crookston & Stage 1999). The OGSI variable is a composite index ranging from 0-100 that is calculated from 4 elements: density of large diameter live trees per hectare, density of large diameter snags per hectare, percentage of downed wood greater than 25 cm in diameter, and an index of tree diameter diversity computed from tree densities in different diameter classes. For live trees and snags, “large diameter” is dependent on forest type and is defined for twelve vegetative zones, each with a unique minimum diameter threshold (50-100 cm for live trees, 50-75 cm for snags). Detailed methods for calculating the OGSI variable can be found in Davis et al. (2015); see Supplemental information (Item S1) for more information on integration of the OGSI variable into our model.

We created a variable called percent pine, which was a the combination of percent total basal area of shore pine (*Pinus contorta*), Jeffery pine (*P. jefferii*), and knobcone pine (*P. attenuata*) from GNN. We modeled this variable because martens have been found with sparse shore pine communities in the Oregon Central Coast population (Eriksson et al. 2019; Linnell et al. 2018), and areas with serpentine soils, which are characterized by sparse tree cover of Jeffery and knobcone pines, stunted tree growth, and dense shrub understories (Harrison et al. 2006; Kruckeberg 1984; Safford et al. 2005). Serpentine soils have been associated with martens in northern California (Slauson et al. 2019b). We visually inspected the serpentine soil layer created by the US Fish and Wildlife Service with the percent pine layer (Schrott & Shinn 2020), confirming overlap inland and inclusion of coastal areas.

Humboldt martens have been associated with dense shrub cover throughout their range (Moriarty et al. 2019; Slauson et al. 2007); salal (*Gautheria shallon*) and evergreen huckleberry (*Vaccinium ovatum*) appear particularly important, as the berries of each occur in Humboldt marten diets and also provide food for marten prey items (Eriksson et al. 2019; Moriarty et al. 2019). We modeled salal and evergreen huckleberry occurrence and combined model outputs to create a shrub cover variable (shrub), which was the sum of the probabilities of species occurrence of both respective shrub rasters (Prevéy et al. 2020a; Prevéy et al. 2020b). We modeled our shrub cover variable using Maxent, version 3.3 (Phillips & Dudík 2008) with detailed methods published elsewhere (Prevéy et al. 2020a; Prevéy et al. 2020b). Briefly, these models used locations from 10 sources, including FIA plots, USDA Forest Service Ecology and Bureau of Land Management plots, iNaturalist, Oregon Flora Project, Consortium of Pacific Northwest Herbaria, and the Global Biodiversity Information Forum. We related locations to contemporary (1981-2010) bioclimatic variables from the AdaptWest project (Wang et al. 2016) to depict the probability of species occurrence (1-100%), or expected habitat quality, for each species (Prevéy et al. 2020a; Prevéy et al. 2020b). The evergreen huckleberry model has not been published previously, but we created it using the same methods as the published salal model (Prevéy et al. 2020a; Prevéy et al. 2020b).

Our final biotic variable (mast) represented hardwood tree and shrub species that produce nuts, seeds, buds, or fruits eaten by wildlife. American marten population numbers appear correlated with mast in hardwood forests of New York (Jensen et al. 2012); similarly, fishers (*Pekania pennanti*) appear associated with mast producing-hardwoods in California (Townsend 2019). It is unclear whether Humboldt martens are similarly associated, but their diet appears dominated by berries and animals that feed on berries and seeds such as passerines and chipmunks (*Tamias sp.*) (Eriksson et al. 2019; Manlick et al. 2019; Slauson & Zielinski 2017). Here, our variable consisted of the percent of basal area within 2016 GNN comprised of tanoak (*Notholithocarpus densiflorus*), giant chinquapin (*Castanopsis chrysophylla*), coastal live oak (*Quercus agrifolia*), canyon live oak (*Q. chrysolepis*), and California bay (*Umbellularia californica*).

### Abiotic variables

We incorporated abiotic variables into our model including those depicting temperature (°C), precipitation (cm), cloud cover (%), coastal proximity, percent slope, and topographic position index. We used 30-year normal PRISM variables of Average Annual Precipitation converted to cm and Maximum Temperature in August available at an 800-m scale (1981-2010, PRISM Climate Group, Oregon State University, http://prism.oregonstate.edu, created 10/17/2019). We explored annual data for temperature (2010-2018), but the available 4 km resolution produced artifacts in the model.

We created a coastal low cloud/fog variable for both the mean and interannual standard deviation of coastal low cloudiness from May through September for 22 years (1996–2017). The 4-km resolution coastal low cloudiness record was satellite derived (NASA/NOAA Geostationary Environmental Satellite Imager measurements) as described by Clemesha et al. (2016). We quantified coastal low cloudiness as the percent of time that low clouds were present relative to the number of corresponding valid half-hourly observations in a 24-hour day. We optimized and validated retrieved data using coastal airports.

We created models with the variable Coastal Proximity, which uses PRISM data and combines coastal proximity and temperature advection influenced by terrain (Daly et al. 2003) modified for the western United States (Daly et al. 2008). We derived percent slope and topographic position index from US Geological Survey digital elevation models. Topographic position index is an indicator of slope position and landform category; it is the difference between the elevation at a single cell and the average elevation of the user-defined radius around that cell (Jenness 2006).

### Scale optimization

We optimized spatial scales for each variable included in our models. We smoothed variables using the extract function in package *raster* in R (Himjmans 2020; R Core Team 2020) with a radius of 50 m, 270 m, 742 m, and 1170 m. Our smallest scale (50 m, 0.81 ha) provided localized conditions matching likely vegetation and location accuracy. We assumed 270 m (20 ha) was approximately the size of a Humboldt marten core use area, similar to optimized scales of vegetation characteristics used in predicting conditions for marten rest structures elsewhere in California (Tweedy et al. 2019). We estimated home ranges for individuals with >30 locations and <400 sq m expected location error (shifted locations were provided in Data S2; Lindenmayer & Scheele 2017). Female Humboldt marten home range estimates averaged 0.8 km^2^ (80 ha) in the Central Coast (n = 6 females, Linnell et al. 2018) and 2.4 km^2^ (260 ha) using a comparable minimum convex polygon estimate in northern California (n = 9 females, PSW 2019). The average of all known telemetered Humboldt marten females was 1.73 km^2^ (173 ha, Table S1) so we used the scale of 742 m (174 ha) as a representation of female home range size. Our broadest scale was based on the largest size of a Humboldt marten male home range (1170 m, 428 ha, Table S1), assuming a male would overlap multiple females and could be interpreted as the smallest unit of population level selection (Linnell et al. 2018; PSW 2019). We used univariate linear models (glm) using our training location data (1) and a random background sample of 9,600 points within the MCP (0) with a binomial distribution. Similar to prior examples (Cushman and McGarigal 2002, McGarigal et al. 2016, Wasserman et al. 2010), we selected the scale for each variable that had the most extreme, and thus the most predictive, coefficient. We visually inspected the fit of each spatial scale using boxplots (Figs. S1, S2, S3).

We visually evaluated whether our final variables were similar between all marten locations, thinned marten locations, surveyed locations without detections (non-detection), and random locations.

### Predicted distribution

We used MaxEnt modeling software v3.4.1 (Phillips et al. 2006) to estimate the probability of Humboldt marten presence within the modeling regions (Merow et al. 2013). MaxEnt uses a machine learning process to develop algorithms that relate environmental conditions at documented species presence locations to that of the surrounding background environment in which they occurred (Elith et al. 2011; Phillips & Dudík 2008). We excluded variables with highly correlated predictors (|Pearson coefficient| > 0.5), selecting the variable that was most interpretable for managers (Table S2). During this process, we considered the variance inflation (Table S3), which allows for evaluation of correlation and multicollinearity. Variance inflation factors equal to 1 are not correlated and factors greater than 5 are highly correlated as determined by (1/(1-R *_i_*^2^)), where R*_i_*^2^ is squared multiple correlation of the variable *i* (Velleman & Welsch 1981).

Within each model iteration, we selected the bootstrap option with 10 replicates, random seed, and 500 iterations. We trained our models using a random subset of 75% of presence locations and tested these using the remaining 25% with logistic output. We used the default of 10,000 random background samples. We varied the response functions to include linear, product, and quadratic features. We selected the “auto features” option for all runs, which allows Maxent to further limit the subset of response features from those selected by retaining only those with some effect.

Maxent provides two metrics to determine importance of input variables in the final model: percent contribution and permutation importance. Percent contribution measures the increase in likelihood associated with modeling rules based on each environmental variable and then divides this by the total gain in likelihood to calculate percent of the total gain associated with each variable (Elith et al. 2011). Maxent keeps track of which environmental variables are contributing to fitting the model, and at each step its algorithm increases the gain of the model by modifying the coefficient for a single feature; the program assigns the increase in the gain to the environmental variable(s) upon which that the feature depends (Phillips 2006). Percent contribution converts these increases to percentages at the end of the training process (Phillips 2006). Halvorsen (2013) produced simulation results suggesting percent contribution can be more informative with uncorrelated environmental variables. This metric is often used to assess variable significance (e.g., Warren et al. 2014).

Permutation importance randomizes the values for each environmental variable between the presences and background points to make that variable uninformative and then measures the resulting drop in area under the curve (AUC; ability to separate known presences from background points). The bigger the drop, the more important that variable was in the overall quality of the model and the resultant data are normalized to percentages (Elith et al. 2011). The permutation importance measure depends only on the final Maxent model, not on the path used to obtain it. Searcy & Shaffer (2016) suggest that permutation importance provides better variable assessment when models and variables are correlated.

Species distribution maps were produced from all models using the maximum training sensitivity plus specificity threshold, which minimizes both false negatives and false positives. We evaluated the AUC statistic to determine model accuracy and fit to the testing data (Fielding & Bell 1997). The AUC statistic is a measure of the model’s predictive accuracy, producing an index value from 0.5 to 1, with values close to 0.5 indicating poor discrimination and a value of 1 indicating perfect predictions (Elith et al. 2006). We assessed variables using response curves, variable contributions, and jackknife tests.

Because over-parameterized models tend to underestimate habitat availability when transferred to a new geography or time period, we used selection methods suggested by Warren & Seifert (2011). Maxent provides the option of reducing overfitting with a regularization multiplier that can be altered by the user to apply a penalty for each term included in the model (β regularization parameter) to prevent overcomplexity or overfitting (Merow et al. 2013; Morales et al. 2017). A higher regularization multiplier will reduce the number of covariates in the model, becoming more lenient with an increased sample size (Merow et al. 2013). We did not include model replicates, an option in the interface, to output the required data (lambda file) and set output to logistic. We altered the Regularization Multiplier from 0.5 to 4 for each 0.5 increment (e.g., Radosavljevic & Anderson (2014).

We ranked candidate models using Akaike’s Information Criterion corrected for small sample sizes (AIC_c;_ Burnham & Anderson 2002). We calculated AIC_c_ using the formula 2k - 2lnL + (2k(k+1))/(n - k – 1). We calculated log likelihood (lnL) by summing the values in all raster cells of the output model, subtracting the log of each cell by the log value of that sum, and extracting and summing the values at the occurrence points (Warren & Seifert 2011). We defined *k* as the number of rows in the lambda files with non-zero lambda values and we considered *n* the number of thinned marten locations. We considered the model with the lowest AIC_c_ value to be the top model, calculated the ΔAIC_c_ value (i.e., change in AIC_c_ value from the top model) for all other models, and considered models with ΔAIC_c_<2 to be competitive models.

For our top model, we generated predicted-to-expected (P/E) ratio curves for our model using only the testing data to evaluate its predictive performance, which was based on the shape of the curves, a continuous Boyce index (Boyce et al. 2002), and Spearman rank statistics. We used the predicted-to-expected curve to inform our suitability thresholds following Hirzel et al. (2006), including predicted unsuitable (P/E and confidence intervals 0-1), marginal (P/E > 1 but overlapping confidence intervals), and suitable (P/E and confidence intervals > 1).

## Results

### Locations

We compiled 1,692 Humboldt marten location records collected between 1996-2020 (baited stations = 542, detection dog = 263, VHF telemetry = 831, roadkill = 15, other = 41 locations). After we spatially-thinned contemporary locations to no more than one per 500 m^2^, 384 locations remained and were generally evenly spread among regions in proportion to their area - Central Coastal Oregon *(n* = 77 locations, 6% of the designated Extant Population Area), Southern Coastal Oregon (*n* = 77 locations, 37% of the EPA), California-Oregon Border (*n* = 33 locations, 3% of the EPA), and Northern Coastal California (*n* = 192 locations, 54% of the EPA) populations, but also included 5 locations that did not occur within the boundaries of these designated populations (USFWS 2019). Location types included den or rest structure locations (18%), genetically-verified scats or telemetry locations (32%), and baited camera or track plate locations (50%).

Our thinned sample had similar medians and data distributions to all contemporary locations with the exception of mast and precipitation, where the medians were slightly lower for our thinned sample. Similarly, surveyed locations without marten detections and random locations were comparable with the most notable different between the medians for salal (Table 1, Fig. 3). Differences between the non-detection and random locations were likely due to clustered sampling efforts (Fig. 1b).

**Figure 2.**
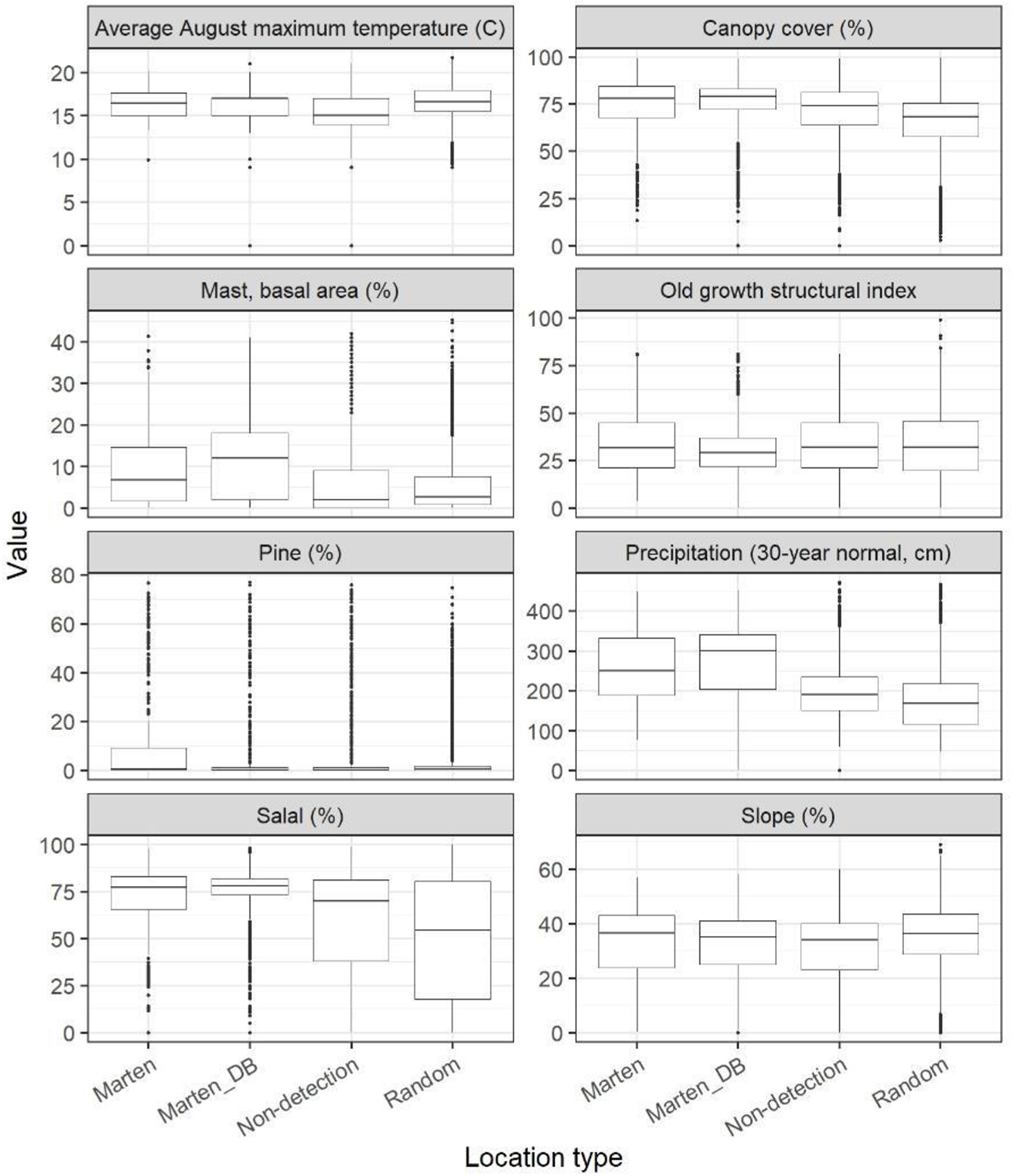
We investigate the range of variables in our thinned dataset compared to all marten locations and detection/non-detection data. To provide the range of values observed in this study, we depict boxplots for the variables in the top model showing the thinned marten data (Marten), all marten known locations (Marten_DB), non-detected but surveyed locations (Non-detection), and random locations within the minimum convex polygon (9,600 random locations).

**Figure 3.**
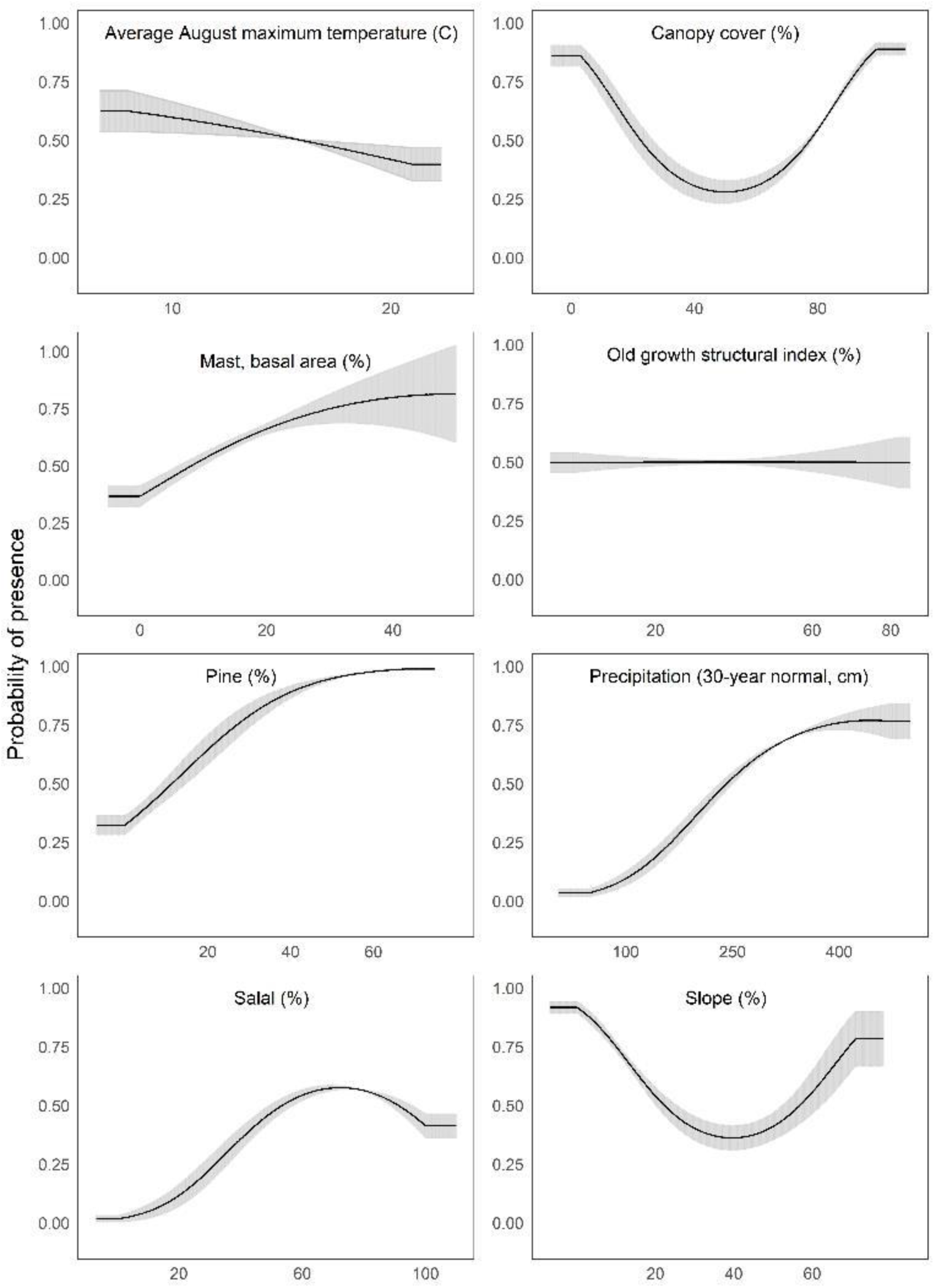
We depict predicted relationships between Humboldt marten locations and each of the variables within our final model. Here, each curve is the predicted probability of presence with no conflicting influence of potentially correlated variables. Humboldt marten locations were correlated with both low and high amounts of canopy cover and percent slope (quadratic response). Predicted distribution was positively correlated with predicted salal (*Gaultheria shallon*) distribution, percentage of pine, precipitation, and mast. We observed a negative correlation between marten locations and August temperature. We observed a slight negative relationship between marten locations and the old growth structural index. Percent contribution and permutation importance values were reported in Table 2. The curves reveal the mean response (black) and standard deviation (gray) for 10 replicate Maxent runs.

**Table 2.** We show the percent contribution and permutation importance from our top Maxent model. We ordered variables by their percent contribution and report the optimized spatial scale (focal radius in meters), the univariate response type, and whether the univariate dependent plots were generally positively or negatively correlated with Humboldt marten (*Martes caurina humboldtensis*) locations.

| Variable | Scale | Response | Univariate Relationship | Percent contribution | Permutation importance |
| --- | --- | --- | --- | --- | --- |
| Salal | 1170 | Quadratic | + | 23.3 | 15.5 |
| Percent pine | 1170 | Product | + | 22.5 | 30.3 |
| Precipitation_30 year average | 1170 | Product | + | 21.6 | 25.3 |
| Canopy cover | 1170 | Quadratic | + | 18.7 | 20.2 |
| Mast | 1170 | Product | + | 5.4 | 1.3 |
| August temperature_30 year average | 1170 | Linear | - | 4.7 | 2.3 |
| Percent slope | 1170 | Quadratic | - | 2.7 | 4.4 |
| Old growth structural index | 50 | Linear | - | 1.2 | 0.7 |

### Distribution modeling

Our model included 8 variables with <60% correlation (Table S2) with variance inflation factors <5 (Table S3). All variables in our model had an optimized spatial scale at our largest extent (1,170 m) except OGSI (50 m), suggesting these variables were most closely matched to martens at the home range scale or fine-scale. Our top model had a Regularization Multiplier of 1.5. The top model, in order of percent contribution, included a positive relationship with salal (23.3%), percent pine (22.5%), average annual precipitation (21.6%), canopy cover (18.7%), and mast (5.4%) followed by a negative relationship with average maximum August temperature (4.7%), percent slope (2.7%), and OGSI (1.1%, Table 2). The permutation importance was similar with the top four variables highly contributing - but with a slightly modified order of percent pine (30.3%), average annual precipitation (25.3%), canopy cover (20.2%), and salal (15.5%) for the top variables (Table 2). The OGSI variable contributed least for both metrics.

We interpreted Maxent’s univariate response curves, but have provided the marginal plots as a supplemental (Fig. S4). Marten locations were correlated with both low and high amounts of canopy cover and percent slope (quadratic response, Fig. 3). We suspect these could be biologically correlated in that extensive flat areas in the Central Coast also have low tree canopy cover, but high shrub density. Moderate amounts of canopy cover (e.g., 5-50%) appeared to be negatively correlated with marten locations. Predicted distribution was positively correlated with predicted salal distribution with some likelihood of a threshold at high values (Fig. 3), percentage of pine (Fig. 3), average annual precipitation (Fig. 3), and mast (Fig. 3). We observed a negative correlation between marten locations and August temperature (Fig. 3). We observed a slight negative relationship between marten locations and OGSI (Fig. 3).

Our final combined model predicted versus expected curve placed unsuitable area <14%, suitable areas of 15-30%, and predicted highly suitable at >30% (Fig. 4) with an AUC value on the test data at 92.1%. The model depicted southern Oregon and northern California as having the largest extent for marten distribution, including areas south of the current known population (Fig. 5).

**Figure 4.**
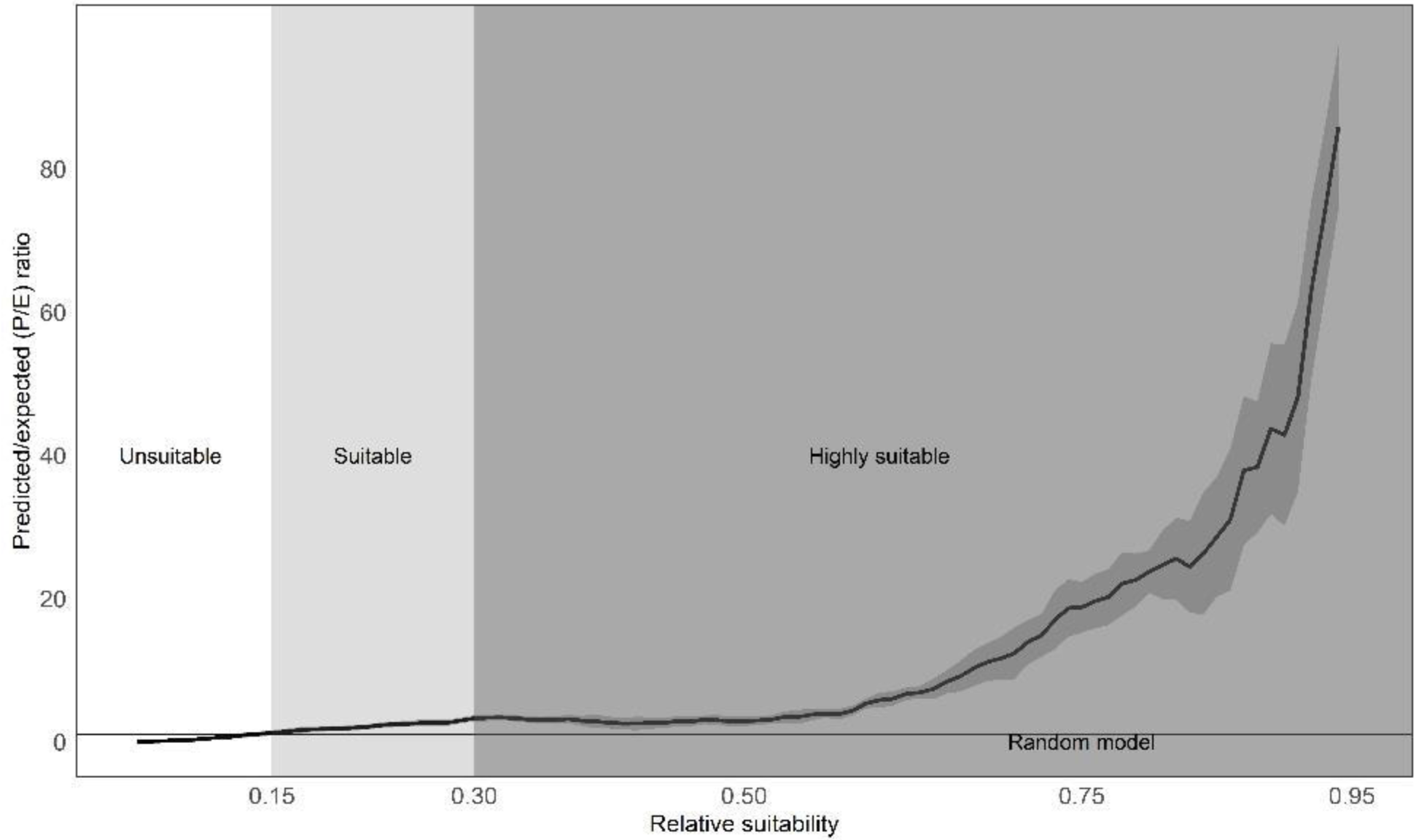
Our predicted suitable transitions for Humboldt marten (*Martes caurina humboldtensis*) range. We present mean predicted vs. expected curve (solid black line) from our model replicates, showing 95-percent confidence intervals (gray-shaded vertical bars). The P/E = 1 threshold is where the curve crosses the random chance line (horizontal orange line), and the blue dashed vertical lines are the 95-percent confidence intervals. We used the predicted-to-expected curve to inform our suitability thresholds following Hirzel et al. (2006), including predicted unsuitable (P/E and confidence intervals 0-1), marginal (P/E > 1 but overlapping confidence intervals), and suitable (P/E and confidence intervals > 1; map depicted in Fig. 5).

**Figure 5.**
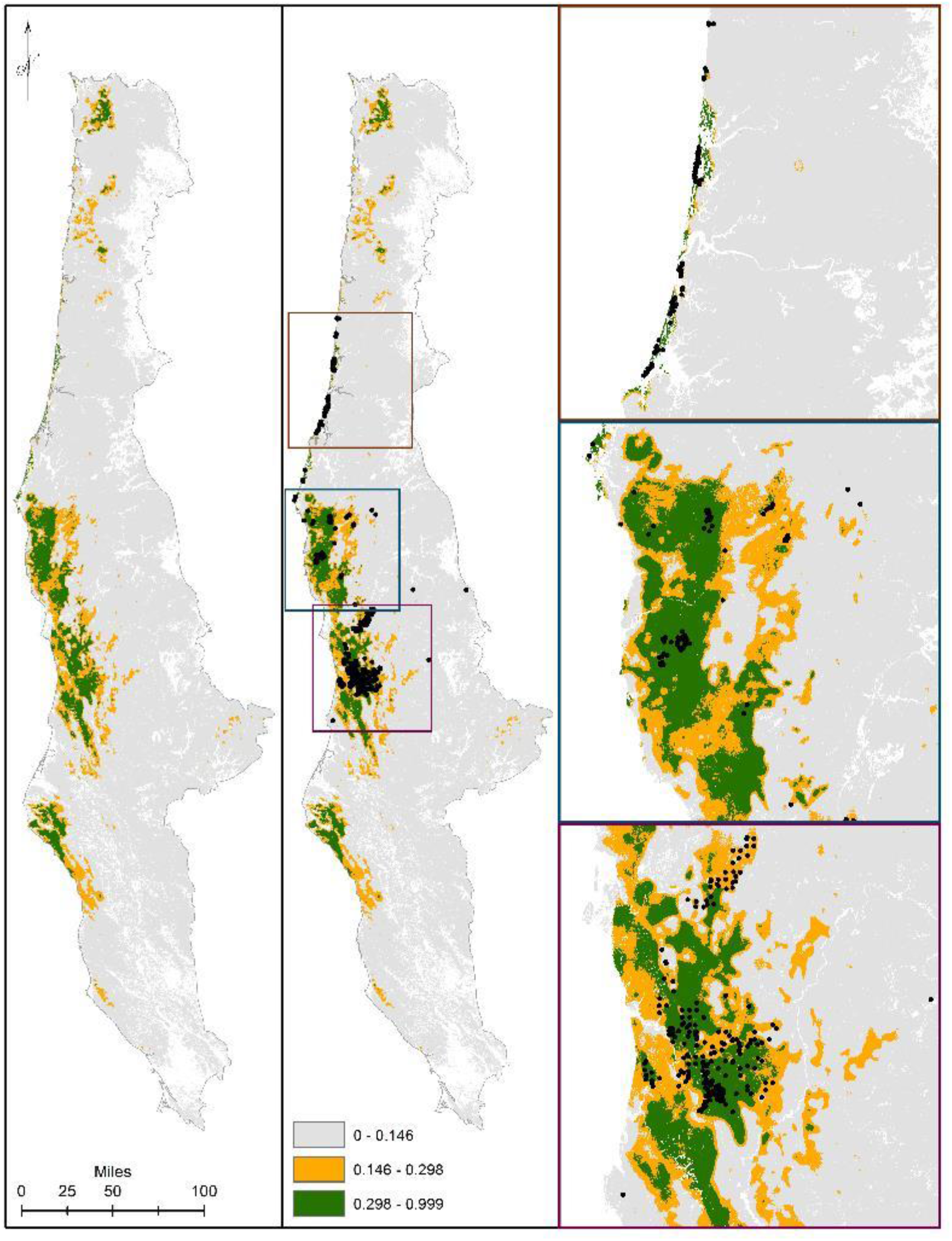
We display our modeled predicted range for Humboldt marten (*Martes caurina humboldtensis*). For predicted range, we followed Hirzel et al. (2006) with predicted versus expected ratios transitioning between predicted highly suitable (green), suitable (orange), and marginal or not predicted suitable (gray). Marten location information was displayed (black dots). We zoomed to population extents to provide increased visual resolution within the Central Oregon Coast (Panel 3a), South coast (Panel 3b), and northern California (Panel 3c).

## Discussion

Our distribution model both predicts areas where Humboldt marten populations are known to occur and identifies areas of potential occurrence outside of known population extents. However, as with all models, there are limitations associated with our predictions, and a clear assessment of these constraints is critical for model results to be accurately used to inform management decisions (Sofaer et al. 2019). For example, abiotic variables such as precipitation or slope are predictive of habitat use but may be uninformative in a management framework because they cannot be altered. Nonetheless, abiotic variables such as temperature and precipitation provide alternative information that can be used for contemporary climatic baselines and future climate scenarios. However, establishing patterns of causality is challenging, and it is also plausible that these abiotic variables are linked to occurrence because remnant Humboldt marten populations happen to occur in wet and steep regions. Further, interpretation of model variables requires consideration of biological (e.g., biotic and abiotic) processes within a spatial context, and to achieve this, we included predictors that provided spatially-accurate depictions of habitat use at the most predictive spatial scale (McGarigal et al. 2016). The culmination of these efforts resulted in a predicted distribution map and hypotheses regarding resources correlated with Humboldt martens.

The role of biotic interactions in shaping the distribution of species has been reported (e.g., Forchhammer et al. 2005, Guisan and Thuiller 2005). However, evidence of the importance of biotic variables alongside abiotic variables for predicting distributions at larger spatial scales has been largely lacking (e.g., Wisz et al. 2013). Here, we show that inclusion of certain abiotic variables in species distribution models can improve model predictions markedly, providing support for the role of biotic interactions in shaping species distributions at the landscape-scale (Wisz et al. 2013). Occurrence of pine and canopy cover were two biotic variables with high model contributions. We interpret the combination of the pine-dominated vegetation type with the quadratic response of canopy cover as copacetic; that is areas with pine dominant forest types typically have lower canopy cover and require adequate access to light for growth, including areas with relatively low canopy cover (Augusto et al. 2003). In these areas, shrub cover, particularly salal and mast-producing shrubs, may be more prevalent. High shrub cover appears to be a consistent component of Humboldt marten habitat in both California (Slauson & Zielinski 2009, Slauson et al. 2007) and Oregon (Moriarty et al. 2019). Predicted marten distribution also was related to increased canopy cover, a finding consistent with other research indicating that martens seek areas with relatively high cover (Bissonette et al. 1997, Hargis et al. 1999). In the univariate responses, canopy cover was quadratic, suggesting a relatively low correspondence with moderate levels of canopy cover. More information is needed to understand if there is an avoidance of thinned forests with moderate levels of canopy cover similar to other Pacific marten literature (Moriarty et al. 2015; Moriarty et al. 2016). Additional information quantifying amounts and extent of shrub and canopy cover are needed to better understand how martens interact with vegetation at fine spatial scales, the combination of low canopy cover with a dense understory layer of shrub and mast producing species or a high level of canopy cover represent achievable conditions that can guide management or target restoration.

Alternatively, biotic variables such as percent pine may not offer similar guidance. Here, they may act as “proxy” variables to conditions where martens currently persist. The influence of percent pine in our predicted Humboldt marten model is rooted in a biological truth, as well-drained or serpentine soils prohibit extensive canopy closure thus promoting extensive shrub growth. However, the presence of pine species is most likely an artifact of co-occurrence with dense shrub cover. Our interpretation concerns are especially salient for historically-occupied areas that no longer share similar attributes, such as previously extensive shrub fields with succession into homogenous but not yet structurally diverse forests.

Biotic variables influencing predicted Humboldt marten distribution in our model were consistent with previous literature with some exceptions, most notably forest age and old growth characteristics. Forest age was not a strong predictor in any model iteration, nor was OGSI. We incorporated OGSI into our model, despite its meager contributions to various model iterations (<5% contribution), because of its high predictive contribution in the Slauson et al. (2019b) model. Within our model, the predicted relationship between Humboldt marten distribution and higher OGSI values was not only weak but negative – higher OGSI values could be interpreted as less suitable for Humboldt marten use (Supplemental Item 1). We suspect the difference between our model and the Slauson et al. (2019) model in regard to OGSI was due to our survey efforts being greatly increased in geographic scope and located martens in a greater diversity of vegetation types with widely differing ages. For example, the Slauson et al. (2019) model included 8 marten locations in Oregon where our model included 167 marten locations with survey efforts randomly or systematically distributed (Moriarty et al. 2018; Moriarty et al. 2019). This resulted in a relatively even proportion of locations throughout the distribution of the Humboldt marten, including telemetry locations in younger forests previously identified as unsuitable (Zielinski et al. 2001). The Slauson et al. (2019b) model had an advantage of using non-detection data and our predictive response curves were similar in both models – both slopes were relatively flat or decreasing with relatively broad confidence intervals. The most obvious difference between models is geographic scope and extent, which is a challenge for non-stationarity.

Although our model indicated that Humboldt marten locations are negatively associated with OGSI, we are not suggesting an avoidance of old forest and characteristics associated with resting, denning or foraging. Rather, OGSI appears to be a poor predictor for Humboldt marten distribution when considering the entire range, perhaps because of a potential mismatch in the association between OGSI and shrub cover. For example, while mature Douglas fir-dominated forests inhabited by Humboldt martens in California may have a substantial shrub component (Slauson et al. 2007), much of coastal Oregon and California is dominated by areas of mature western hemlock, a shade-tolerant species that is strongly associated with reduced shrub cover (Kerns & Ohmann 2004). When examining the individual contributions of OGSI and marten locations in a model only with the components of OGSI, downed wood was the most influential variable (Supplemental Item S1). In other words, Humboldt marten distribution appears to be more strongly associated with shrub cover and understory conditions rather than moderately old forests with a density of larger trees or snags – with the mismatch being that some areas where Humboldt martens occur (e.g., mature Douglas fir forest; Slauson et al. 2007) are characterized by both older forest conditions and substantial shrub cover, while other areas (e.g., serpentine or coastal pine forests; Eriksson et al. 2019, Moriarty et al. 2019) are characterized by substantial shrub cover, but not older forest conditions. Contemporarily, throughout the range, occurrence does not appear to be correlated with higher OGSI values, which suggests a potential overemphasis of the OGSI variable in the Slauson et al. (2019) model – particularly in Oregon with few marten locations to base inference. Applications of the Slauson model to Oregon was an extrapolation beyond the range of the data (Romesburg 1981; Rosenbaum 2010).

Both models have challenges in interpretation from non-stationary effects. Non-stationarity in this context describes species-habitat relationships that are variable across time or space and can fundamentally alter interpretation of model output. Here, we were unable to account for temporal non-stationarity (e.g., winter, summer) because the location data seldom had locations surveyed in both seasons. We might expect habitat selection to be more critical in the summer when martens are rearing kits and mating, similar to findings in Martin et al. (In press), but we could not evaluate such influence. It would be deeply flawed to assume that our model, and any other range-wide model, could accurately address the local variation of predictive variables between populations at fine spatial scales. Similarly, models are based on data collected to as of summer 2020 and we suspect there are more martens east in the Oregon coast (e.g., in the vicinity of Ashland, Applegate based on new locations) but our model would not predict such areas due to sparse location information. Future endeavors could develop site-specific models, ideally using telemetry data that are biologically linked with fitness (e.g., long-lived adult female rest and den structures) to address predicted habitat. Understanding habitat was not our goal with this exercise; nonetheless, our modeling exercise provides opportunities to test the influence of these predictive covariates with independent data.

Abiotic factors such as increased precipitation, proximity to the coast, and cool temperatures could be an indication of vegetation growth or may be indicative of a physiological temperature threshold. Precipitation, an unalterable condition but potentially a factor that may be altered in a changing climate, was one of the top 3 predictive variables, fulfilling a high proportion of the model contribution (>20%) in all model simulations. Range limit theorems have long postulated the importance of elevation, altitude, and weather establishing the upper limits of species distributions (e.g., Darwin 1859). If these variables are causally linked to marten occurrence, a plausible mechanism is that cooler wetter conditions are related to increased dense vegetation, which probably aids martens in avoiding predators. Predation was documented as the single greatest limitation to a small and endangered fisher population (Sweitzer et al. 2015) and is a plausible hypothesis for Humboldt marten distribution limitation. Coupled with additional food from berries and small animals that eat berries (e.g., birds, rodents), these areas may create exceptional, yet typically uncharacteristic, habitat (Eriksson et al. 2019). A second synergistic mechanism aligns with Humboldt marten persistence in areas with relatively stable climates (e.g., a backwards velocity evaluation of past change) and areas that may remain relatively stable in future climate predictions (Belote et al. 2018; Carroll et al. 2015). Evidence for this hypothesis would align with observations of martens associated with cooler temperatures associated with higher precipitation within their range limits. The global distributions of “boreal” martens (*Martes caurina, M. americana, M. martes, M. zibellina, M. melampus*) are north of 45 degrees latitude, and these species reside in areas with deep snow cover for a large portion of the year (Krohn et al. 2004; Suffice et al. 2020). The thermoneutral zone of martens has yet to be explored. Generally, species range limits appear highly influenced by abiotic factors (e.g., Normand et al. 2009), although there have been several situations where biotic factors such as vegetation, competition, or predation appear to influence distributions (Louthan et al. 2015; Sirén & Morelli 2020). Here, we would predict that the stable cool and wet climate in near-coastal forests may provide Humboldt martens with a less energetically-expensive temperature gradient.

This modeling effort can be integrated into the framework proposed in the Humboldt marten conservation assessment and strategy, which described a 3-prong approach for Humboldt marten conservation: (1) protect existing populations; (2) re-establish populations in areas with habitat; and (3) restore or focus management efforts to improve habitat (Slauson et al. 2019a). A fundamental foundation of protecting existing populations includes identifying population locations and geographic extents, describing population sizes, and monitoring changes in populations over time. Our model predicts habitat outside of the currently understood boundaries of Humboldt marten populations; prioritizing survey effort in areas that are predicted to be habitat but are of unknown marten occupancy would be an important first step in protecting existing populations. A stratified-random approach, expending survey effort in both predicted suitable and unsuitable areas, would best evaluate marten occurrence. Incorporating non-invasive survey methods (e.g., remote cameras), coupled with genetic analyses, further presents an opportunity to effectively survey species that are cryptic and occur at low densities over large spatial and temporal scales, and to robustly answer questions concerning animal distribution, abundance and demographics (Pauli et al. 2010, Lamb et al. 2019).

Consideration of areas where Humboldt martens do not appear to occur in contemporary times, but our model predicts as suitable habitat, is exemplary of interpretation that requires careful evaluation of biological processes. For example, our model predicts marten distribution in the northern and southern extremes of our modeling region, including the Tillamook State Forest in northern Oregon coast and the King Range National Conservation Area in California, respectively. While the Tillamook State Forest had higher amounts of the abiotic variables of our model (e.g., high precipitation amounts, cool temperatures, and coastal proximity), there were fewer areas predicted with forest vegetation conditions there (Supplemental File 1). Alternatively, the suitability of the unoccupied southern modeling extent (King Range National Conservation Area) was primarily driven by biotic factors (e.g., high canopy cover, high amount of masting trees and shrubs; Supplemental Fig. x). Given the importance of these biotic factors in our model, and that the southern portion represents a significant portion of the historical range of the Humboldt marten (Zielinski et al. 2001), it may provide marten habitat that is currently unoccupied. While suitability as proxy for population viability in a given area has been questioned broadly (Stephens et al. 2015), potential but unoccupied habitat should be areas considered for future survey efforts.

While our model predicts habitat in apparently unoccupied areas that could eventually be considered for population re-establishment and restoration, it also underscored our lack of reliable knowledge on critical and consistent components of Humboldt marten habitat, and we suggest it is premature to pursue actions such as population re-establishment without first refining our understanding of marten habitat requirements. This effort and prior explorations of vegetation associations exemplify the vast range of vegetation conditions used by Humboldt martens (Eriksson et al. 2019; Moriarty et al. 2019; Slauson et al. 2007). Yet while some vegetation conditions are certainly required for species’ population persistence, vegetation conditions may not act as a categorical surrogate for habitat.

Although species’ distributions can be influenced by very apparent biotic (e.g., vegetative) and abiotic (e.g., temperature thresholds) conditions, they may also be strongly influenced by less-apparent factors such as interspecific interactions with predator or competitor species (Siren 2020). As an example, spotted owls (*Strix occidentalis*) closely align with old-growth forest conditions which have been characterized with relatively high accuracy (Davis et al. 2016), yet spotted owl population viability is dramatically decreased with presence of barred owls (*S. varia*) due to competition, predation, and predation risk (Diller et al. 2016; Dugger et al. 2016; Wiens et al. 2014). Although few examples exist for carnivores, a recent evaluation suggests that while lynx (*Lynx lynx*) distributions are closely-tied to deep snow, the influence of reducing bobcat (*L. rufus*) competition was stronger than the influence of snow itself (Siren 2020). A directed research effort would be necessary to understand the relative importance of vegetation structures, vegetation types, prey, predation, and competition for Humboldt marten persistence.

We developed a range-wide species distribution model based on extensive survey coverage and modern vegetation and climatic variables. Although these results provide predictions for habitat components, describing habitat would be best informed by measures of survival and fecundity rates. At present, managers have conflicting sources of evidence that can lead to confusion in interpretating of Humboldt marten ecology. In this paper, we provided models that are complementary, but not similar to other models (Schrott & Shinn 2020; Slauson et al. 2019b), particularly in regard to interpretation of OGSI. Instead of interpreting differences between models as a conflict, we posit this as evidence of the conservation challenge described by Caughley (1994) and in our bison example. We lack enough information regarding where Humboldt martens resided historically to compare with our contemporary distribution (Loehle 2020), and we are generally ignorant of population densities, causal associations of population declines, and population limitations. Such an understanding is essential to describe expectations of future range (Brown et al. 1996). Each available case study provides an imperfect piece of evidence, and the lack of consistency among studies is suggestive of imperfect knowledge of what components constitute Humboldt marten habitat. To avoid “wicked problems” from differing views for rare species conservation (e.g., Gutiérrez 2020; Jones et al. 2020), amassing information collaboratively with a goal of prospective meta-analyses and study-level replication will be essential (Facka & Moriarty 2017; Nichols et al. 2019).

### Conclusions

Based on our modeling and an evaluation of available evidence, we conclude that the most consistent range-wide characteristic with Humboldt marten distributions are vegetation associations with extensive dense shrub cover or complex understory vegetation, which may ultimately be an association with increased food availability or predation escape cover. An understanding of the strength of these interactions and factors that limit populations will ultimately be needed for making wise conservation decisions. Moving forward with conservation decisions with incomplete knowledge necessitates an adaptive management framework.

## Supporting information

Supplemental Item S1

## Acknowledgements

The desire and decision to request an updated model incorporating newer presence and habitat information was from the Oregon Humboldt Marten stakeholder group, which is facilitated by the U.S. Fish and Wildlife Service in Oregon. Non-invasive marten surveys were conducted by Pacific Northwest and Southwest Research Stations, Oregon State University, Humboldt State University, Green Diamond Resource Company, NCASI, the Siuslaw, Rogue-Siskiyou, and Six Rivers National Forests, Hancock Forest Management, Weyerhaeuser, Oregon Department of Forestry, and the Confederated Tribes of Siletz Indians of Oregon. Detection dog surveys were completed by Rogue Detection Dog Teams and the former group within Conservation Canines, University of Washington. Considerable aid with field logistics, vehicles, housing, and equipment was provided by the U.S. Fish and Wildlife Service, Salem District BLM, USFS Rogue River-Siskiyou and Siuslaw National Forests, Weyerhaeuser, Hancock Forest Management, and USFS Region 6 Regional Office. We obtained private land access or surveys were completed by trained staff within the ownership for all randomly selected survey points – thanks to Weyerhaeuser, Hancock Forest Management, Starker Forests, and Roseburg Timber for access or data. Reviews by Drs. D. Miller and J. Verschuyl, B. Hollen, our anonymous peer reviewers, and the Associate Editor improved previous versions of this manuscript. Extreme thanks to all field crew leaders (S. Smythe, M. Linnell, B. Peterson, G. W. Watts, J. Bakke, C. Shafer, K. Kooi, and M. Penk) and team members (E. Anderson, D. Baumsteiger, A. Benn, J. Buskirk, B. Carniello, M. Cokeley, S. Hart, P. Iacano, A. Kornak, T. McFadden, E. Morrison, A. Palmer, T. Peltier, N. Palazzotto, S. Roon, S. Riutzel, C. Scott, K. Smith, R. Smith, T. Stinson, M. Williams, B. Woodruff, and K. Wright). Any use of trade, firm, or product names is for descriptive purposes only and does not imply endorsement by the U.S. Government.

## Supplemental Information Captions

**Supplemental Data S1. We provide marten and random locations with associated modeled values at multiple scales.**

We provide raw data from thinned Humboldt marten (*Martes caurina humboldtensis*) and random locations within the known area (minimum convex polygon). We extracted data at 4 spatial scales (radius = 50m, 270m, 742m 1170m) associated with Humboldt marten biology (e.g., Table S1) and ran univariate general linear models to select the most predictive spatial scale.

**Supplemental Data S2. We provide marten locations from two radio telemetry studies for the purpose of estimating home range size.**

We provide raw data from Humboldt marten (*Martes caurina humboldtensis*) locations collected within the Northern California (VHF telemetry) or Oregon Central Coast (GPS telemetry) regions with methods described elsewhere (Delheimer et al. In press; Linnell et al. 2018). We used only data from individuals with greater than 30 locations and with location error estimated at less than 400 sq m to estimate range size (Table S1). We used range size to inform spatial scale smoothing extents for modeled variables. Because Humboldt martens are state endangered in California and federally threatened, we were not comfortable revealing location data (Lindenmayer & Scheele 2017). For each marten, we selected either added (heads) or subtracted (tails) the individual marten’s location by a random value (range = 50,000 – 100,000 m) such that all location data for a marten were similar spatially for the purpose of estimating range size but not relatable to the true location where the marten was monitored.

**Supplemental Item S1. Exploration of the Old Growth Structural Index and Humboldt martens (*Martes caurina humboldtensis*)**

Because the old growth structural index has been interpreted as a variable indicative of Humboldt marten predicted locations in other models, but not within our exploration, we explore the variable in detail and explain nuances to why there might be differences.

**Supplemental Figure S1, S2. We display boxplots for all considered biotic variables with predicted associations with Humboldt martens.**

To provide the range of values observed in this study, we depict boxplots for the biotic variables in the top model showing the thinned marten data and random locations (25/marten location; 9,600 random locations) for each spatial scale (radius = 50m, 270m, 742m 1170m) associated with Humboldt marten biology (e.g., Table S1, described in methods).

**Supplemental Figure S3. We display boxplots for all considered abiotic variables with predicted associations with Humboldt martens.**

To provide the range of values observed in this study, we depict boxplots for the abiotic variables in the top model showing the thinned marten data and random locations (25/marten location; 9,600 random locations) for each spatial scale (radius = 50m, 270m, 742m 1170m) associated with Humboldt marten biology (e.g., Table S1, described in methods).

**Supplemental Figure S4. We depict predicted relationships (marginal plots) between Humboldt marten locations and each of the variables within our final model.**

Here, each curve is the predicted probability of presence given the other variable responses. Predicted distribution of Humboldt martens were correlated with increasing canopy cover, percent pine, precipitation, temperature, mast, percent salal (*Gaultheria shallon*) distribution, and the old growth structural index. We observed a negative correlation between marten locations percent slope. Percent contribution and permutation importance values were reported in Table 2. The curves reveal the mean response (black) and one standard deviation (gray) for 10 replicate Maxent runs.

**Supplemental Table S1. Humboldt marten home range sizes by sex and region.** We compiled home range size information for Humboldt martens in the Central Coast of Oregon and Northern California to base our scale optimization for variables in this distribution model. Data from northern California were extracted from unpublished data (PSW 2019) and data from the Central Oregon Coast were in the supplemental material within Linnell et al. (2018).

**Supplemental Table S2. We evaluated Pearson correlation coefficients for inclusion in our distribution model.**

We evaluated Pearson correlation coefficients and restricted variables with correlation >0.6 to guide variables in a Humboldt marten (*Martes caurina humboldtensis*) distribution model. For our final model, we selected canopy cover over the diameter diversity index. We selected OGSI over tree age or diameter diversity index due to its use in prior Humboldt marten range wide models. We selected salal over huckleberry because of salal’s presumed structural use (Eriksson et al. 2019, Moriarty et al. 2019).

**Supplemental Table S3. We evaluated Variance Inflation Factors, which reveal correlation and multicollinearity, to guide variables in a Humboldt marten (*Martes caurina humboldtensis*) distribution model.**

The absolute value of variance inflation factors equal to 1 are considered not correlated and values greater than 5 are highly correlated as determined by Velleman & Welsch (1981). Here, diameter diversity index conflicted with canopy cover and we used canopy cover in our final model for the ease of interpretation and use in a management context.

