## Supplemental Item S1 for "Predicted distribution of a rare and understudied forest carnivore: Humboldt martens (*Martes caurina humboldtensis*)"

### **Supplemental Item S1. Investigation of the variable “old growth structural index” as a surrogate for Humboldt marten habitat. An exploration of inconsistencies between two range-wide habitat models.**

### Supplemental item for the manuscript:

### Predicted distribution of a rare and understudied forest carnivore: Humboldt martens (*Martes caurina humboldtensis*)

### Katie Moriarty^1^, Joel Thompson^2^, Matthew Delheimer^3^, Brent Barry^4^, Mark Linnell^5^, Taal Levi^6^, Keith Hamm^7^, Desiree Early^7^, Holly Gamblin^8^, Micaela Szykman Gunther^8^, Jordan Ellison^1^, Janet S. Prevéy^9^, Jennifer Hartman^10^, Ray Davis^11^

Here, we provide a detailed evaluation and comparison of the results of our range-wide Humboldt marten habitat model, presented within this manuscript, with a previously-published model (Slauson et al. 2019). In particular, the Slauson et al. (2019) model places substantial emphasis on Humboldt marten occurrence being strongly and positively associated with an “old growth structural index” variable (hereafter, OGSI); however, we found little evidence for a similar relationship within our model. Given that OGSI is already being used as a “surrogate” for Humboldt marten habitat (e.g., Schrott and Shinn 2020, Supplemental Item Fig. 1), it may behoove managers and wildlife practitioners to understand the differences between variables incorporated into our respective models and their influences on subsequent model outputs.

***What is the old growth structural index and how is it calculated?***

OGSI is a composite index that combines several spatially-explicit, remotely-derived forest structure elements using Lemma’s gradient nearest neighbor index (Ohmann and Gregory 2002). OGSI was designed to describe the continuum of forest succession, with higher values in the later stages of succession (Spies and Franklin 1988). OGSI extends in geography to Washington, Oregon, and California and was created in part to monitor old forest conditions over broad spatial scales (Davis et al. 2015), especially areas within the Northwest Forest Plan and over the range of the Northern Spotted Owl (Davis et al. 2016).

As an index, OGSI has evolved in both complexity and precision over time. For example, the 2006 version of OGSI was calculated from 5 elements, including: tree age (age_dom), density of large live trees (>100 cm in diameter; tph_ge_100cm dbh), a diameter diversity index computed from tree densities in different diameter classes (ddi), density of large snags (stph_5015), and percentage of downed wood greater than 25cm in diameter (dvph_ge_25). The 2006 version of OGSI had the same inputs for all vegetative zones in the Pacific Northwest (see code block below for more detail) and index values ranged between 0 and 100. Since 2010, OGSI has been calculated from 4 elements: density of large live trees per hectare (ltphc), density of large snags per hectare (stph_ge), percentage of downed wood greater than 25cm in diameter (dcov_ge_25cm), and an index of diversity of tree diameter computed from tree densities in different diameter classes (ddi). Unlike the 2006 OGSI version, the more recent version has twelve vegetative zones that each have a unique threshold for what is considered a “large” live tree or a “large” snag ranging from 50-100cm for live trees and 50-75cm for calculating snag densities. In other words, this metric is dependent on forest type – for example, a “large diameter” tree or snag in a lodgepole pine (*Pinus contorta*) stand would be comparatively considered “small in diameter” within a coastal redwood (*Sequoia sempervirens*) stand. The index ranges between 0 and 100 see Davis et al. (2015) and the infographic on page 6.

***Why use the old growth structural index?***

With too much data or too few replicates, condensing variables into a composite index, such as OGSI, can be a useful tool for modeling. There are both formal and practical procedures for creating such indices. Formal methods are often applied, for instance, if a goal is to describe vegetation associations with hundreds of variables (e.g., canopy cover, number and diameter of each tree species, stems per shrub, leaves per shrub), as a model would be computationally intractable with too many variables. In wildlife, common opportunities to reduce many variables into 2 or 3 composite variables include principal component, generalized discriminant, and canonical correlation analyses (Ramsey and Schafer 2002). Similarly, but less formally, one can combine correlated variables with *a priori* hypotheses or biological logic by adding the values. The challenge of interpreting such data is that the results are an index and not a feature. For instance, instead of describing the relationship of Humboldt marten locations to canopy cover, one would describe the relationship between “axis 1” and “axis 2” or an index without defining which components are most related to the species of interest.

The challenge of index interpretation and relating to biological expectations is not unique to OGSI. Similarly and more simply in wildlife habitat relationships, quadradic mean diameter (QMD) is $\sqrt{(\sum d_{i}^{2})/n}$ where *d* is the diameter of an individual (i) tree at breast height and n is the number of trees (Curtis and Marshall 2000). Quadratic mean diameter has been used in silviculture since the early 1900’s (e.g., Graves 1908) and is one of the primary components in California’s Wildlife Habitat Relation database to assign a habitat value (e.g., high or low quality) to a location based on vegetation elements (Salwasser and Laudenslayer 1982, Garrison 1994). By the nature of the calculation, the QMD value of a given forest stand increases when it is thinned, as the result of the removal of small diameter trees (Curtis and Marshall 2000). While habitat quality is generally presumed to improve with increasing QMD values for many forest-dependent species such as Pacific martens, processes such as forest thinning may in fact degrade habitat quality (e.g., Stephens et al. 2014, Moriarty et al. 2016), despite the appearance of improvement (i.e., increased QMD). As such, interpretation of indices such as OGSI or QMD can be challenging and not associated with biological realities if the situational components are not clearly described.

***Humboldt marten locations and OGSI***

We initially modeled using the 2016 version of the OGSI variable, despite its meager contributions to our model iterations (<5% contribution), primarily for purposes of comparison with the Slauson et al. (2019) model. When incorporated into our model, the relationship between OGSI and Humboldt marten locations was not only weak but also often negative – higher OGSI values could be interpreted as less suitable for predicted Humboldt marten use.

We assume that both modeling efforts used the best available data, but the breadth of surveys for Humboldt martens have exponentially increased since 2015. The Slauson et al. (2019) model used data collected from 1996-2010. Data were mostly from northern California, but there were 8 locations in Oregon (compared to 48% of our data), and the majority of locations used within the Slauson model were collected at the spatial scale for fishers (~6 km spacing) during other efforts (e.g., Carroll et al. 1999, Zielinski et al. 2010), which means the spacing may have missed the martens with their smaller home ranges and rigid territoriality (Moriarty et al. 2017). Although the text describes 1,159 considered surveys, the model used 559 non-detection and 44 detection locations (Table 5) with detection data strongly being spatially autocorrelated (e.g., Figure 5; Slauson et al. 2019).

During our study, we also combined efforts from various Humboldt marten focused studies and study designs (Slauson et al. 2007, Barry 2018, Linnell et al. 2018, Moriarty et al. 2018, Gamblin 2019, Moriarty et al. 2019), and we included the detection data used in the previous model. We prioritized marten rest and den locations as well as scat locations. Our marten surveys in Oregon were randomly or evenly distributed throughout the entire coast range, including all forested age classes (Moriarty et al. 2018, Moriarty et al. 2019). We included an even proportion of locations throughout the range, including telemetry data in younger forests previously identified as unsuitable (Zielinski et al. 2001). We compiled 5,153 locations (1,692 detections, 3,461 non-detections) from 1996-2020, thinned the data to one location within a 500-m by 500m cell, and modeled based on 384 locations (see associated manuscript for details).

Although named the same, the way OGSI was represented differs between the Slauson et al. (2019) model and both our effort and Schrott and Shinn (2020) due to how the metric was calculated. Slauson et al. (2019) summed the cumulative 2006 OGSI score within each spatial scale. For the final model, the variable used at the 1-km radius for spatial scale, provided a range of observed locations from an observed index between 0 to 8,000 (Slauson et al. 2019b; Fig. 3). In contrast, Shrott and Shinn (2020) used 2012 GNN (LEMMA 2014) but averaging a 977m radius area and creating a “habitat core” from OGSI values >36 and areas >1500ha. Here, the index value of 36 was selected because it was the median for values within the depicted historical range - not because there was any tested association with Humboldt marten locations (pg. 30; Schrott and Shinn 2020; Supplemental Item Fig. 2).

We investigated these assumptions using summary data and models. A histogram of marten location data was not immediately suggestive that there were more marten locations with increased OGSI values (Supplemental Item Fig. S3). Similarly, our thinned Humboldt marten locations were not extremely different than random locations at any measured spatial scale (Supplemental Item Fig. S4). We calculated OGSI and each component (forest age, diameter diversity index, large snag density, large tree density, and downed wood). We used the same modeling procedure described within our manuscript but only included OGSI or the 5 components. Our model with OGSI as the sole variable to evaluate Humboldt marten distribution performed similar to a random variable (Supplemental Item Fig. S5). Our Humboldt marten model with each of the 5 OGSI components did better in creating a more interpretable map, likely because the response curves could vary (Supplemental Item Fig. S6). Here, the variables that explained the most variation were percentage of downed wood and the diameter diversity index (Supplemental Item Table S1).

Regardless of study design and variable calculations, we suggest Slauson et al. (2019) may have overemphasized the importance of OGSI and underestimated the implications of the abiotic factors. Within Slauson et al. (2019) the top-ranked model in included 4 variables: (1) Old-growth structural index (OGSI) at a 1km scale; (2) Serpentine at 3km scale; (3) Precipitation; and (4) Adjusted elevation. The biotic variable, OGSI, was then adopted as a primary variable for a landscape connectivity model (Schrott and Shinn 2020). Interestingly, the partial dependence plot in the Slauson et al. (2019, Figure 3) model appears generally negative with a broad confidence interval, then declines steeply with larger values of OGSI. Similar to the functional response curve (Slauson et al. 2019, Figure 3), we observed a generally neutral or negative relationship of Humboldt marten locations and OGSI with our univariate response curves.

Although Humboldt marten locations appear weakly and potentially negatively associated with OGSI, we are not suggesting that Humboldt martens avoid older structures with complexities such as cavities or mistletoe (Slauson and Zielinski 2009, Tweedy et al. 2019). Such individual structures and microsites such as cavities are strongly linked to resting and denning in Pacific martens and fishers (e.g., Matthews et al. 2019, Tweedy et al. 2019). Nonetheless, vegetation characteristics collected as “plot level” or in the general vicinity may not be predictive of habitat for carnivores such as fishers (Green 2017, Green et al. 2019) because of their landscape scale and fine scale needs. As we describe in our main text, a biological prediction for why OGSI would be a poor predictor for range-wide modeling is because it appears that Humboldt martens are often positively associated with shrubs, but much of coastal Oregon and California have a large occurrence of western hemlock, a shade-tolerant species that is strongly associated with reduced shrub cover (Kerns and Ohmann 2004) but in areas that often have relatively high OGSI values with increased diameter diversity. Further, remotely sensed vegetation data are not necessarily accurate at a pixel by pixel level – evaluations of large live trees and snags using similar vegetation layers suggest such data are appropriate regionally but not at fine spatial scales (Bell et al. 2021). There may be potential to identify structures and predicted habitat using fine-scale LiDAR data (Wing et al. 2015, Joyce et al. 2019), but such data are not yet available range wide. Once additional location data (e.g., telemetry locations) and actively sensed data are combined, we expect an updated predictive range model would be more accurate.

In summary, Humboldt marten locations and predicted habitat are variable in relation to vegetation characteristics like OGSI. Regionally in northern California, variables like OGSI may be correlated to Humboldt marten locations. Nonetheless, based on available data, it is inappropriate to use OGSI as a surrogate for predicted habitat throughout the Humboldt marten range.

Habitat models are an evolving opportunity to learn. We applaud efforts to continue data collection and efforts to address challenging information gaps aimed to inform conversation efforts.

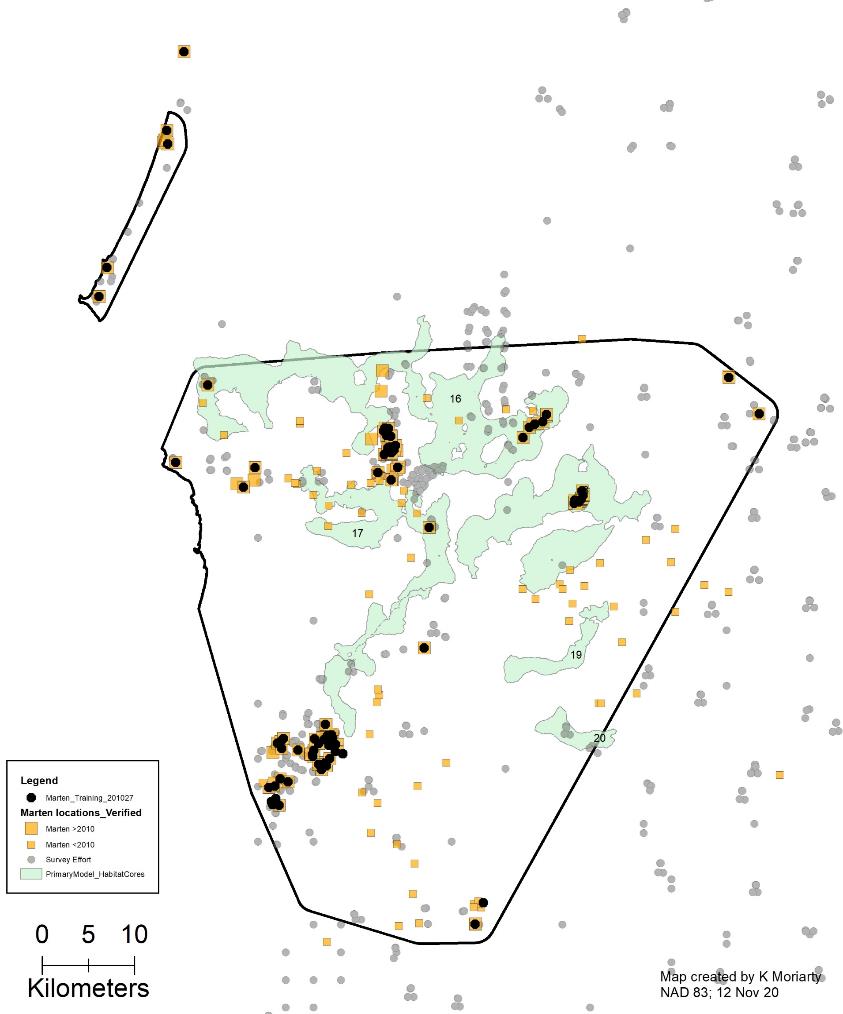

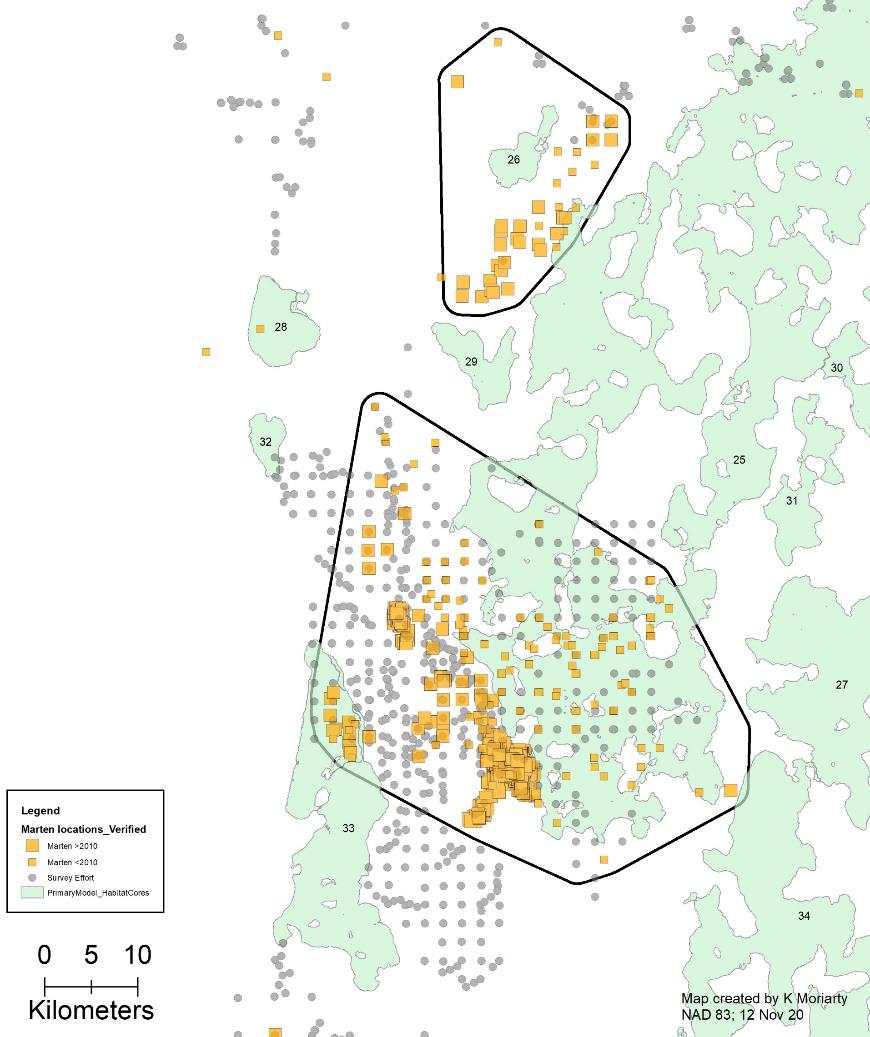

Supplemental Item Figure S1. Prior models use a remotely sensed variable, old growth structural index (OGSI), to depict “habitat cores” (Schrott and Shinn 2020). Here, we provide examples of those cores with survey detection/non-detection locations focused on Humboldt marten distribution. Black lines are areas Humboldt marten population designations, orange are marten locations, grey are surveyed areas. Green polygons are modeled habitat cores (Schrott and Shinn 2020), containing ~9% of known Humboldt marten locations.

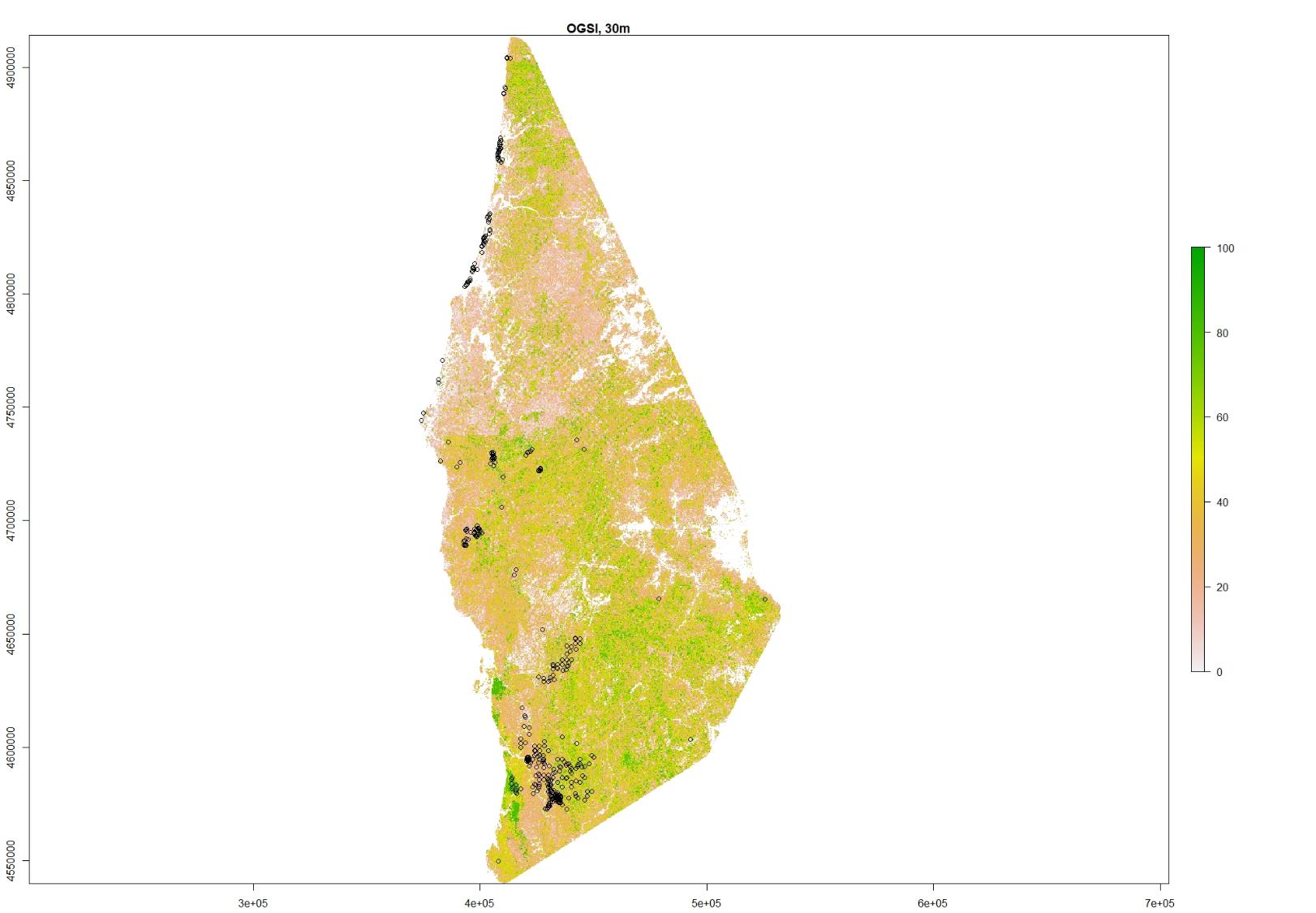

Supplemental Item Figure S2. We display the 2016 remotely sensed index old growth structural index (OGSI) distribution within the current extent of Humboldt marten locations. High values of OGSI are green. Humboldt marten locations are black outlined dots.

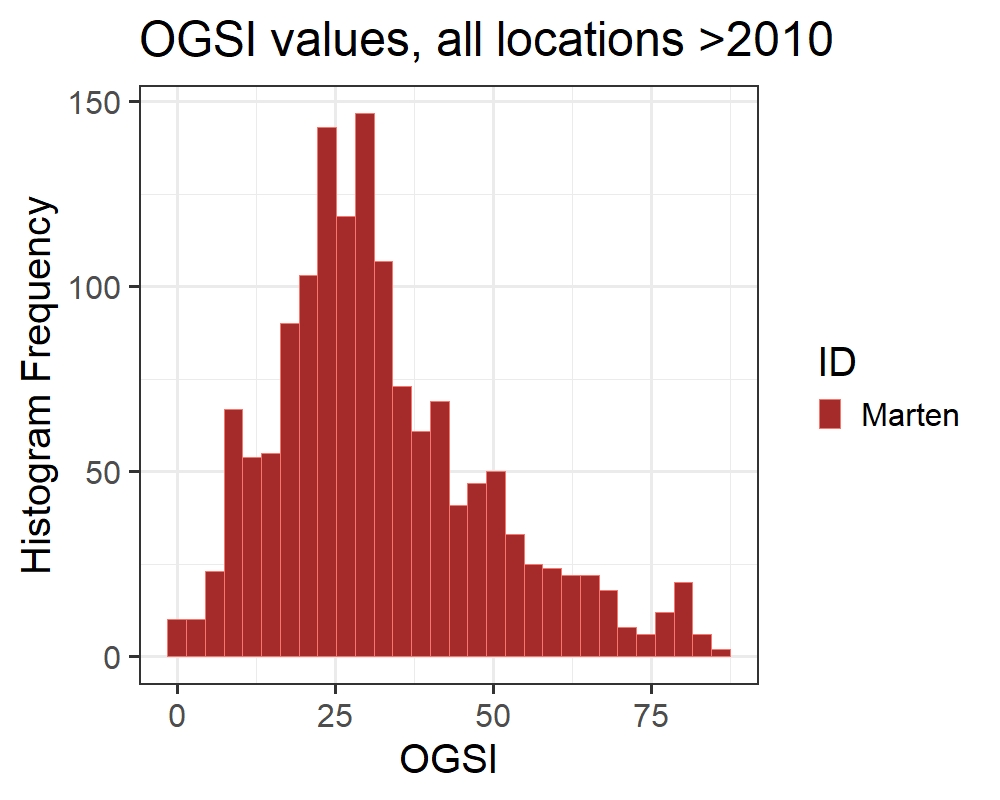

Supplemental Item Figure S3. Histogram of the remotely sensed index old growth structural index (OGSI) and the value for all known Humboldt marten locations. The median value of OGSI within the historic Humboldt marten range with the 2012 vegetation layer was an index of 36 (Schrott and Shinn 2020).

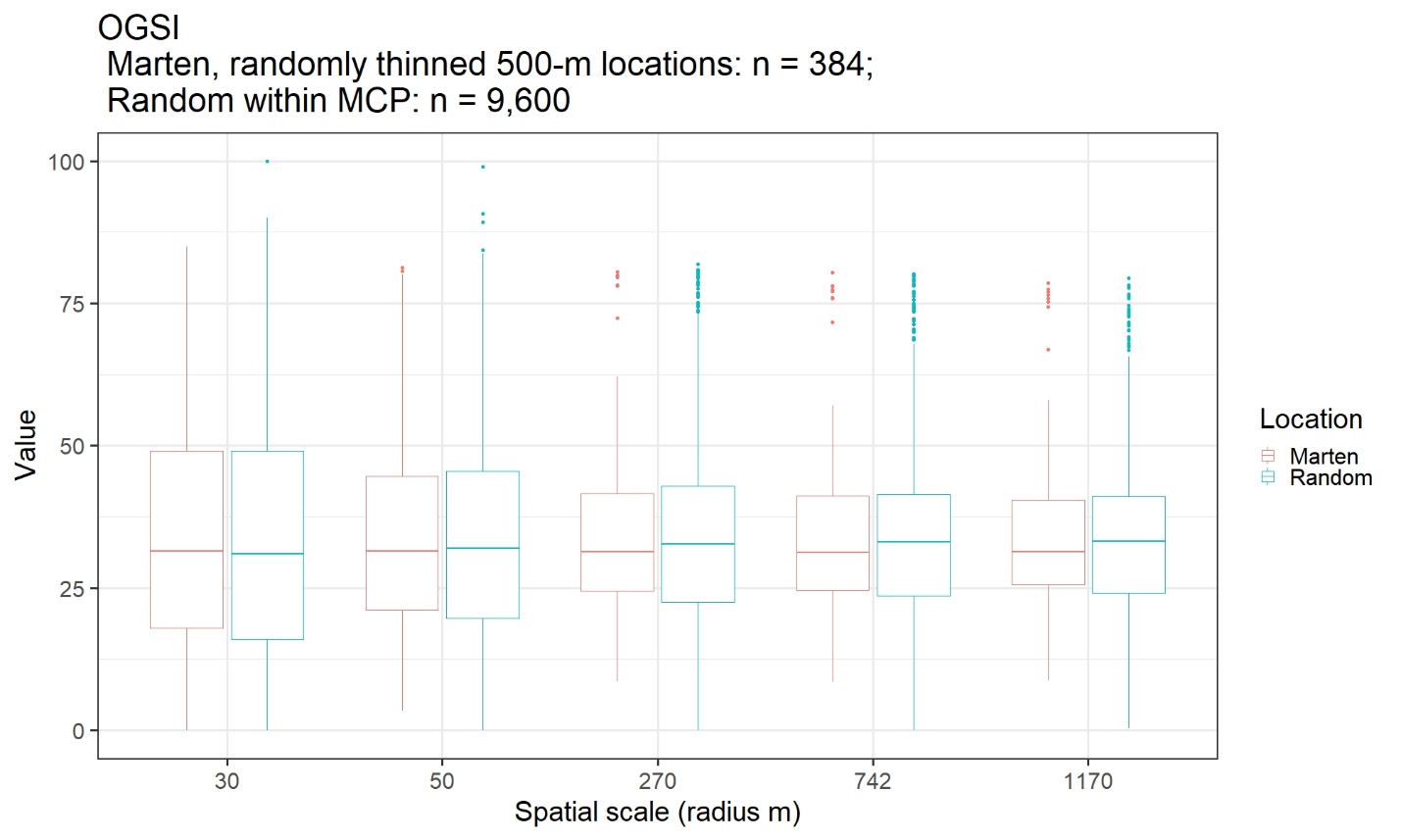

Supplemental Item Figure S4. We compared the spatially thinned location data with 25 random locations per known (9,600) at spatial scales presumed relevant to Humboldt marten biology. Here, note the median for Humboldt marten locations and random locations are similar at each spatial scale. With a focal radius >30m, the median for random values is higher than marten locations. Univariate generalized linear model beta coefficients using these data starting at 50m were 0.0014, 0.00028, -0.00094, and -0.00249 respectively. At large spatial scales (742m, 1170m) the relationship between marten locations and OGSI were slightly negative.

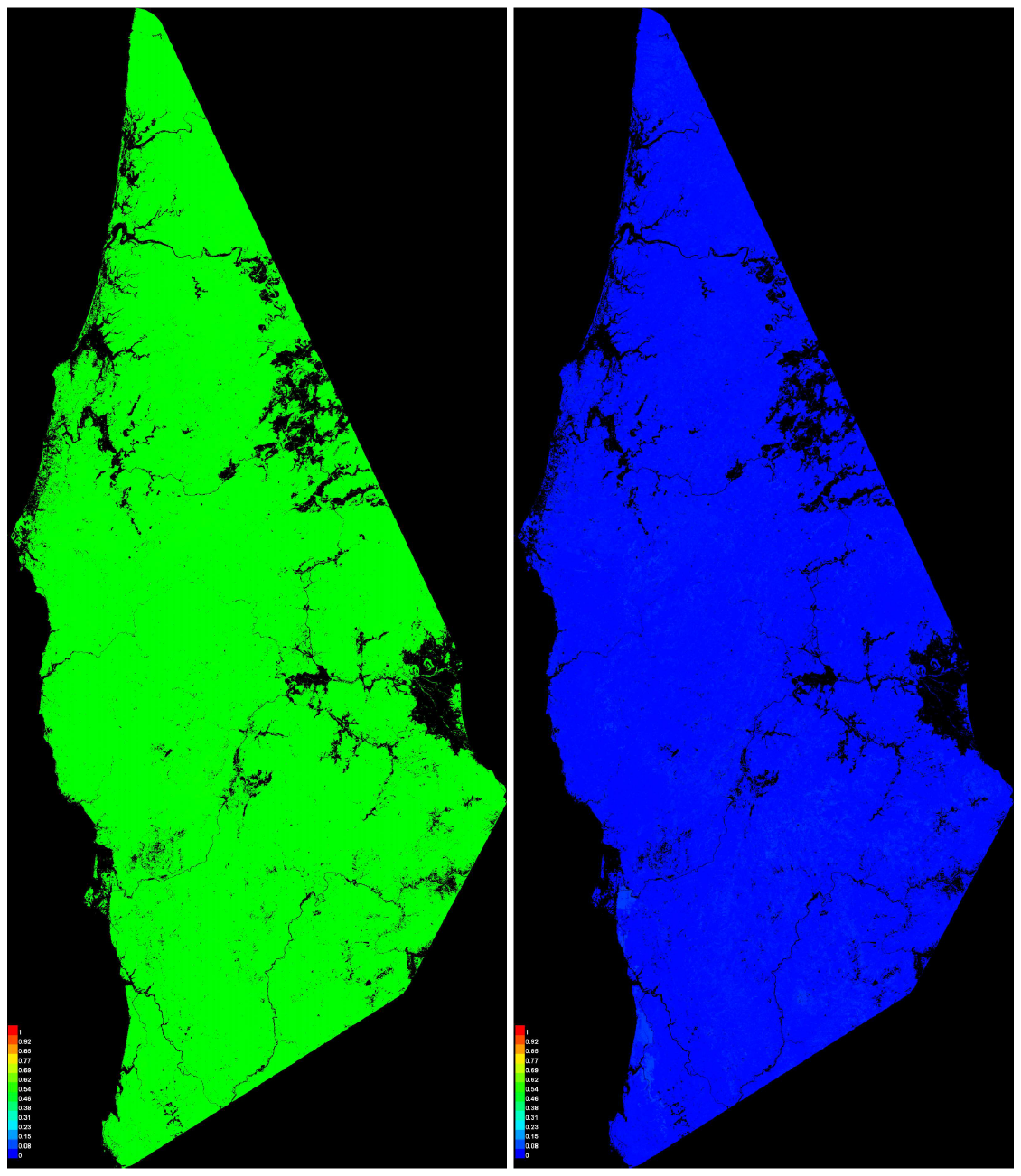

Supplemental Item Figure S5. We created a Maxent model only with the variable old growth structural index (OGSI). Here, it predicted Humboldt marten distribution slightly above a random value. Green is approximately 50% predicted probability and red would indicate high correlation with Humboldt marten locations.

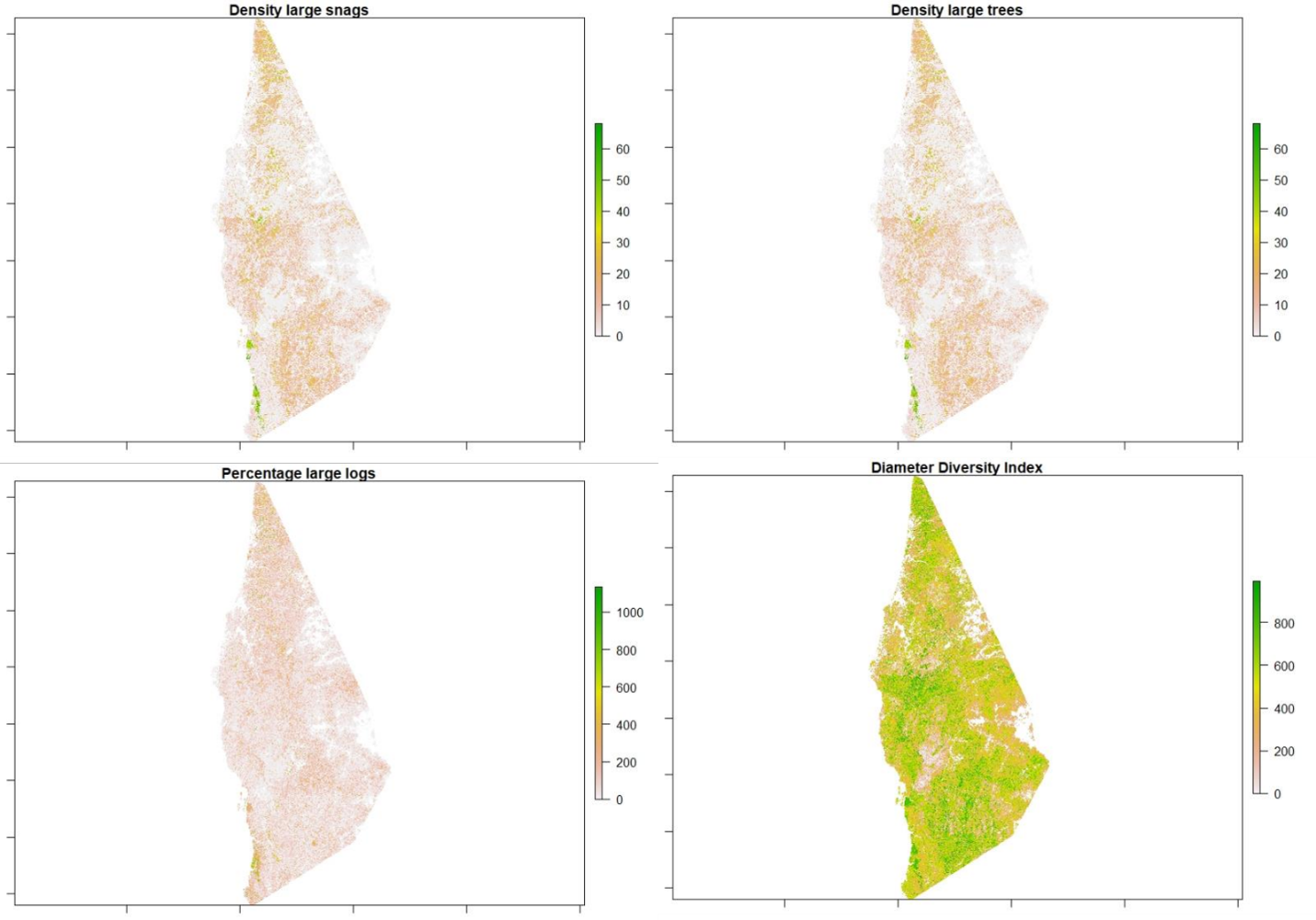

Supplemental Item Figure S6. We separated the index OGSI to each of its components similar to the 2006 GNN product to investigate which element(s) within the OGSI index were correlated with Humboldt martens.

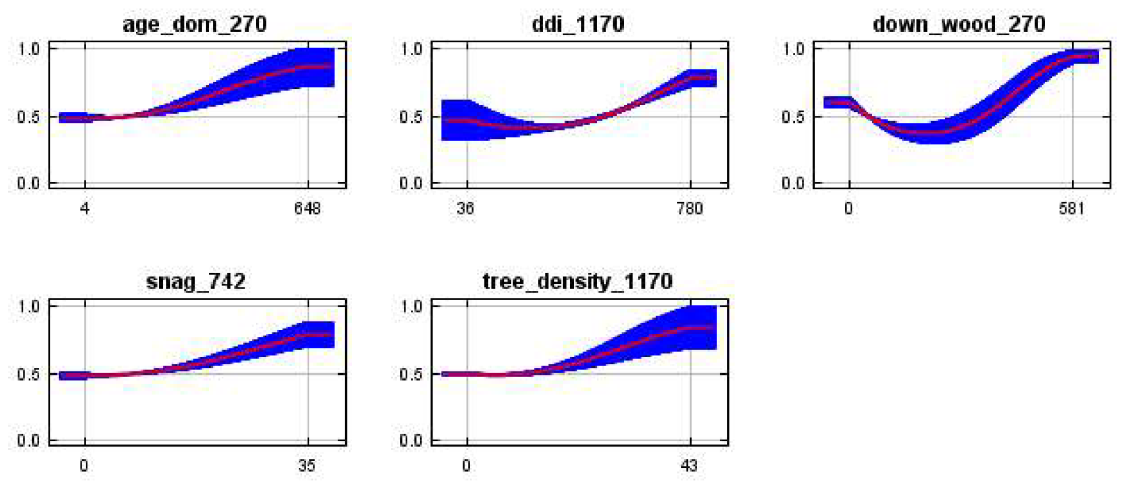

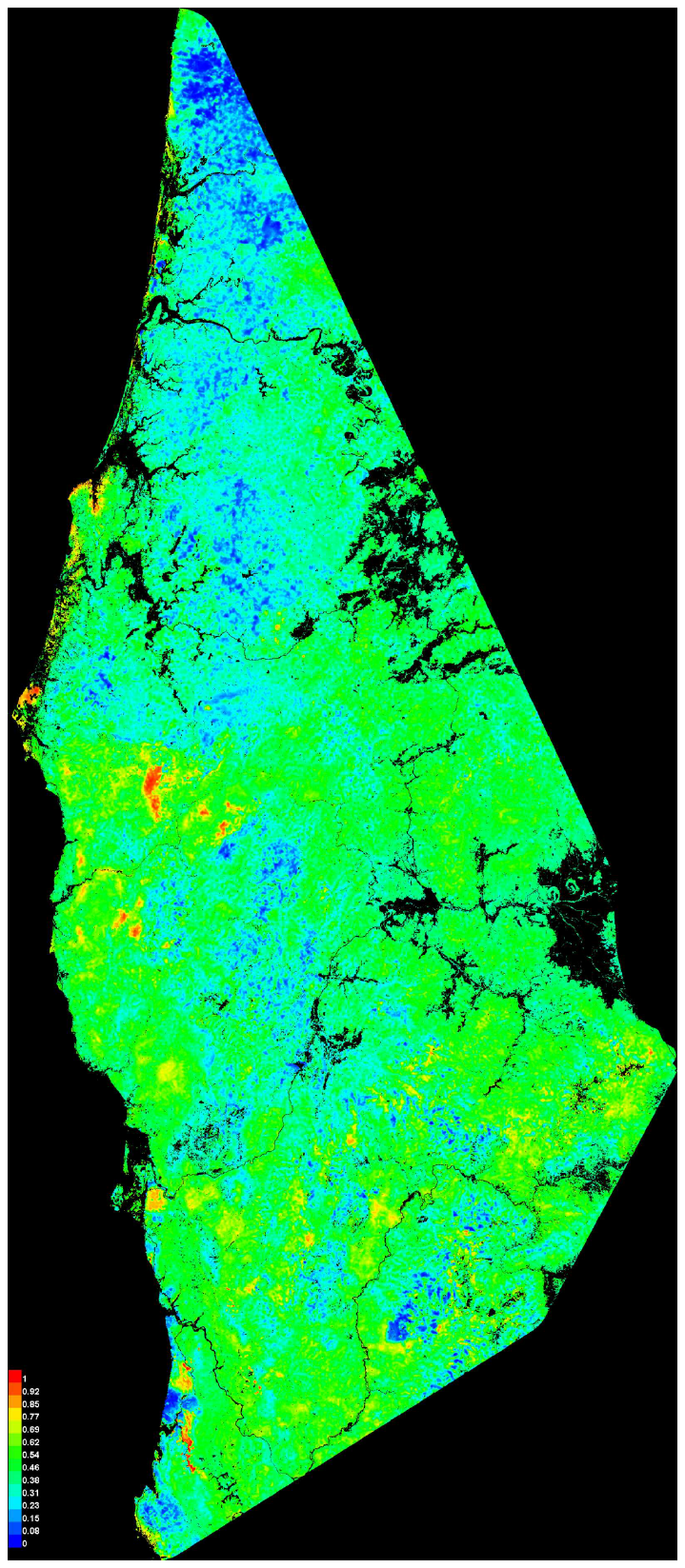

Supplemental Item Figure S7. We depict a spatial map of predicted Humboldt marten range from a Maxent model using the 5 components of the variable old growth structural index (OGSI), with our known and thinned Humboldt marten occurrences (n = 384). From these components, percentage of downed wood at a smoothed radius of 270m (down_wood_270), diameter diversity index at a smoothed radius of 1170m scale (ddi_1170), large tree density (tree_density_1170), large snag density (snag_742) and estimated tree age (age_dom_270) were the order of model rank by percent contribution.

Supplemental Item Table S1. We created a Maxent model using the 5 components of the variable old growth structural index (OGSI). When evaluating either percent contribution or permutation importance, estimated tree age contributed least and either downed wood or diameter diversity contributed most to the predicted model.

| Variable | Scale | Relationship | Percent contribution | Permutation importance |
| --- | --- | --- | --- | --- |
| Downed wood | 270 | + | 36 | 21.2 |
| Diameter diversity index | 1170 | + | 23.5 | 36.2 |
| Large tree density | 1170 | + | 19.2 | 15.9 |
| Large snag density | 742 | + | 13.7 | 14.5 |
| Age dominant forest | 270 | + | 7.5 | 12.2 |

Code for 2006 OGSI index:

CREATE FUNCTION dbo.GET_OGSI

(@age_dom DECIMAL(9,4), @tph_ge_100 DECIMAL(9,4),

@ddi DECIMAL(9,4), @stph_5015 DECIMAL(9,4), @dvph_ge_25 DECIMAL(9,4))

RETURNS DECIMAL(9,4) AS

BEGIN

DECLARE @age_score FLOAT, @tph_score FLOAT

DECLARE @ddi_score FLOAT, @snag_score FLOAT

DECLARE @cwd_score FLOAT, @ogsi DECIMAL(9,4)

--Live tree age

IF @age_dom <= 200.0

SET @age_score = 0.004 * @age_dom

ELSE IF @age_dom > 200.0 AND @age_dom <= 450.0

SET @age_score = 0.64 + (0.0008 * @age_dom)

ELSE IF @age_dom > 450

SET @age_score = 1.0

--Live TPH

IF @tph_ge_100 <= 17.0

SET @tph_score = 0.02941 * @tph_ge_100

ELSE IF @tph_ge_100 > 17.0 AND @tph_ge_100 <= 32.0

SET @tph_score = 0.21667 + (0.01667 * @tph_ge_100)

ELSE IF @tph_ge_100 > 32.0 AND @tph_ge_100 <= 55.0

SET @tph_score = 0.40217 + (0.01087 * @tph_ge_100)

ELSE

SET @tph_score = 1.0

--Diameter diversity index

SET @ddi_score = 0.1 * @ddi

--Snag TPH

IF @stph_5015 <= 1.0

SET @snag_score = 0.5 * @stph_5015

ELSE IF @stph_5015 > 1.0 AND @stph_5015 <= 3.0

SET @snag_score = 0.375 + (0.125 * @stph_5015)

ELSE IF @stph_5015 > 3.0 AND @stph_5015 <= 14.0

SET @snag_score = 0.68182 + (0.02273 * @stph_5015)

ELSE

SET @snag_score = 1.0

--Coarse woody debris volume

IF @dvph_ge_25 <= 40.0

SET @cwd_score = 0.0125 * @dvph_ge_25

ELSE IF @dvph_ge_25 > 40.0 AND @dvph_ge_25 <= 260.0

SET @cwd_score = 0.45455 + (0.00114 * @dvph_ge_25)

ELSE IF @dvph_ge_25 > 260.0 AND @dvph_ge_25 <= 630.0

SET @cwd_score = 0.57432 + (0.00067568 * @dvph_ge_25)

ELSE

SET @cwd_score = 1.0

--Composite old growth habitat index

SET @ogsi =

((@age_score + @tph_score + @ddi_score + @snag_score + @cwd_score)/5.0)*100.0

RETURN @ogsi

END
