## Supplemental Table S1 for "Predicted distribution of a rare and understudied forest carnivore: Humboldt martens (*Martes caurina humboldtensis*)"

**Humboldt marten home range sizes by sex and region.**

We compiled home range size information for Humboldt martens in the Central Coast of Oregon and Northern California to base our scale optimization for variables in this distribution model. Data from northern California were extracted from unpublished data (PSW 2019) and data from the Central Oregon Coast were in the supplemental material within Linnell et al. (2018).

| Marten | Study Area | Estimator | Sex | Age | Area_km2 |
| --- | --- | --- | --- | --- | --- |
| F01 | Northern California | 100% MCP | Female | Adult | 1.4 |
| F02 | Northern California | 100% MCP | Female | Juvenile | - |
| F03 | Northern California | 100% MCP | Female | Juvenile | 3.1 |
| F04 | Northern California | 100% MCP | Female | Subadult | 2.6 |
| F05 | Northern California | 100% MCP | Female | Subadult | 2.6 |
| F06 | Northern California | 100% MCP | Female | Adult | 5.9 |
| F07 | Northern California | 100% MCP | Female | Juvenile | - |
| F08 | Northern California | 100% MCP | Female | Adult | 1.7 |
| F09 | Northern California | 100% MCP | Female | Juvenile | 1.1 |
| F10 | Northern California | 100% MCP | Female | Juvenile | - |
| F11 | Northern California | 100% MCP | Female | Adult | - |
| F12 | Northern California | 100% MCP | Female | Juvenile | - |
| F13 | Northern California | 100% MCP | Female | Subadult | 2.6 |
| F14 | Northern California | 100% MCP | Female | Juvenile | - |
| F15 | Northern California | 100% MCP | Female | Adult | 0.8 |
| M01 | Northern California | 100% MCP | Male | Juvenile | 2.3 |
| M02 | Northern California | 100% MCP | Male | Juvenile | - |
| M03 | Northern California | 100% MCP | Male | Juvenile | - |
| M04 | Northern California | 100% MCP | Male | Subadult | - |
| M05 | Northern California | 100% MCP | Male | Juvenile | - |
| M06 | Northern California | 100% MCP | Male | Subadult | 2.5 |
| M07 | Northern California | 100% MCP | Male | Subadult | - |
| M08 | Northern California | 100% MCP | Male | Subadult | 4.3 |
| M09 | Northern California | 100% MCP | Male | Juvenile | 7.8 |
| M11 | Northern California | 100% MCP | Male | Juvenile | - |
| M12 | Northern California | 100% MCP | Male | Juvenile | - |
| M13 | Northern California | 100% MCP | Male | Subadult | - |
| M14 | Northern California | 100% MCP | Male | Subadult | - |
| M15 | Northern California | 100% MCP | Male | Juvenile | - |
| M16 | Northern California | 100% MCP | Male | Juvenile | - |
| M17 | Northern California | 100% MCP | Male | Juvenile | - |
| M18 | Northern California | 100% MCP | Male | Juvenile | - |
| M19 | Northern California | 100% MCP | Male | Adult | - |
| F01 | Central Oregon | 99% LoCoH | Female | Adult | 0.59 |
| F03 | Central Oregon | 99% LoCoH | Female | Adult | 0.62 |
| F04 | Central Oregon | 99% LoCoH | Female | Adult | 0.79 |
| F05 | Central Oregon | 99% LoCoH | Female | Adult | 0.71 |
| F06 | Central Oregon | 99% LoCoH | Female | Adult | - |
| F07 | Central Oregon | 99% LoCoH | Female | Adult | 0.84 |
| F08 | Central Oregon | 99% LoCoH | Female | Adult | 0.64 |
| M01 | Central Oregon | 99% LoCoH | Male | Adult | 2.2 |
| M02 | Central Oregon | 99% LoCoH | Male | Adult | 2.2 |
| M03 | Central Oregon | 99% LoCoH | Male | Adult | 1.7 |
| M04 | Central Oregon | 99% LoCoH | Male | Adult | 1 |
| Average Female |  |  |  |  | 1.732667 |
| Max Male |  |  |  |  | 4.3 |

Linnell MA, Moriarty K, Green DS, and Levi T. 2018. Density and population viability of coastal marten: a rare and geographically isolated small carnivore. *PeerJ* 6:e4530 - '4521 pg. 10.7717/peerj.4530

PSW. 2019. Humbodlt marten data summary and report for the northern coastal California population: distribution and population parameter estimates. USDI Fish and Wildlife Service, USDA Forest Service Pacific Southwest Research Station. p 33.
