## Supplementary figures and images for "Predicted distribution of a rare and understudied forest carnivore: Humboldt martens (*Martes caurina humboldtensis*)"

### Supplemental Figure S1

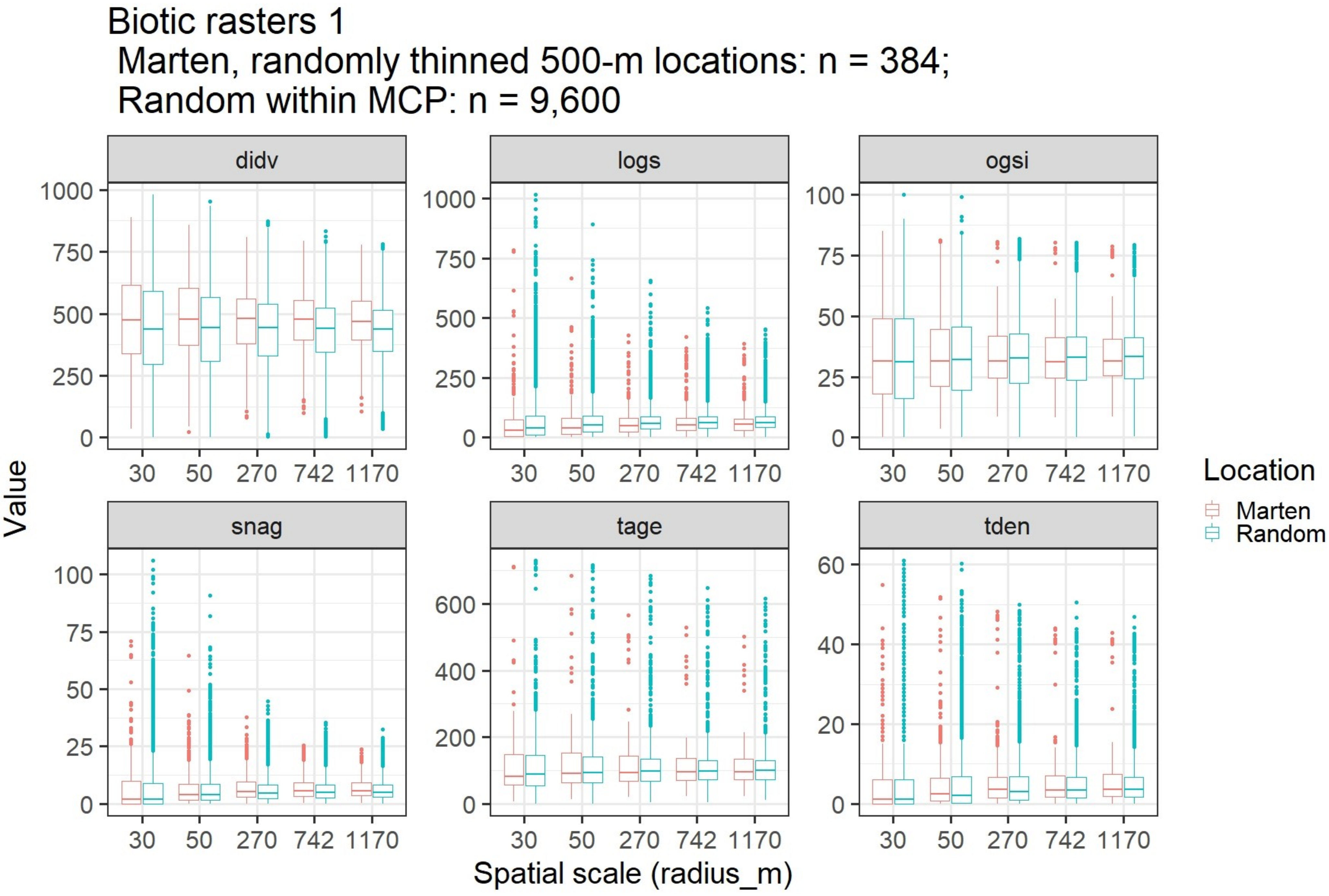

### Supplemental Figure S2

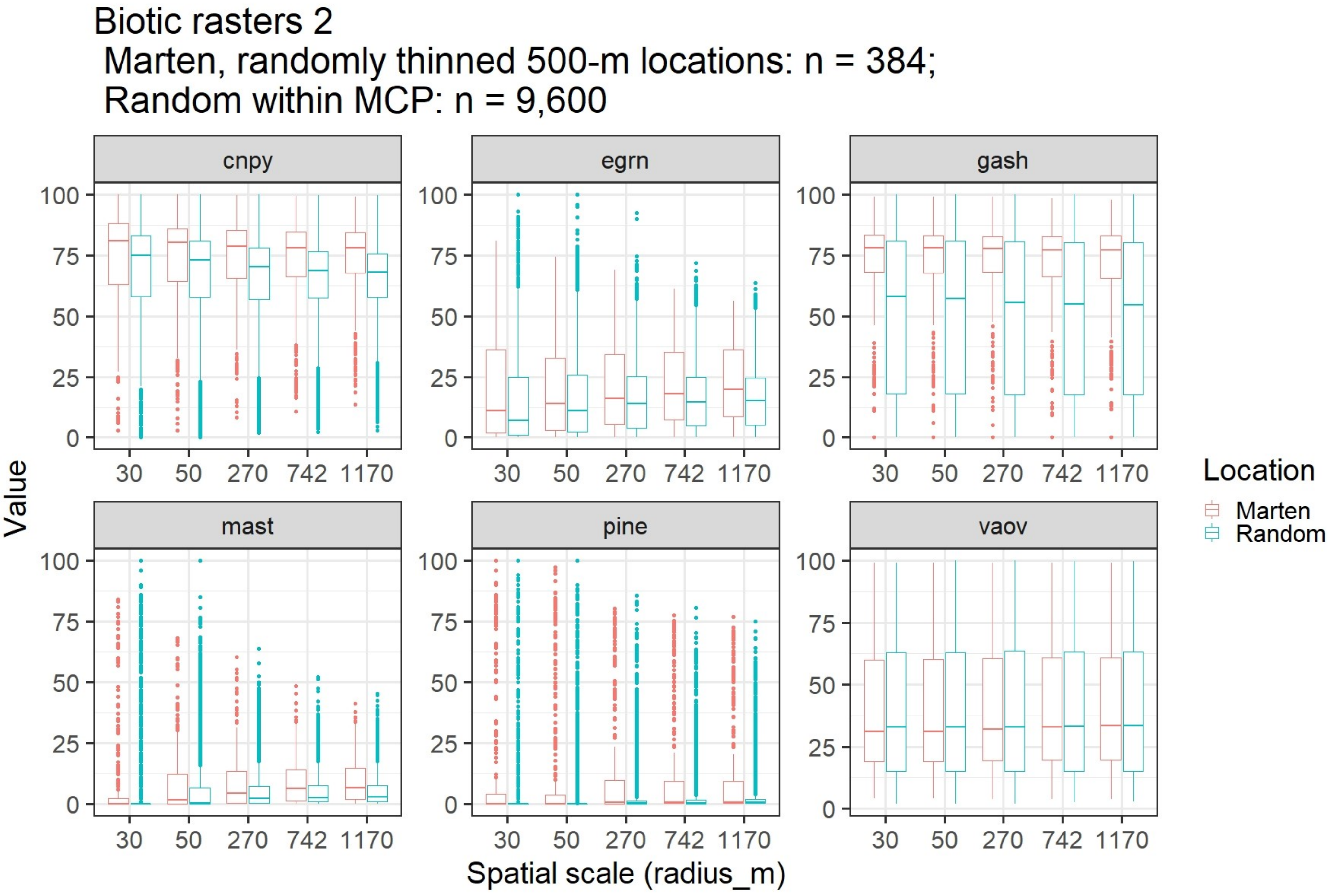

### Supplemental Figure S3

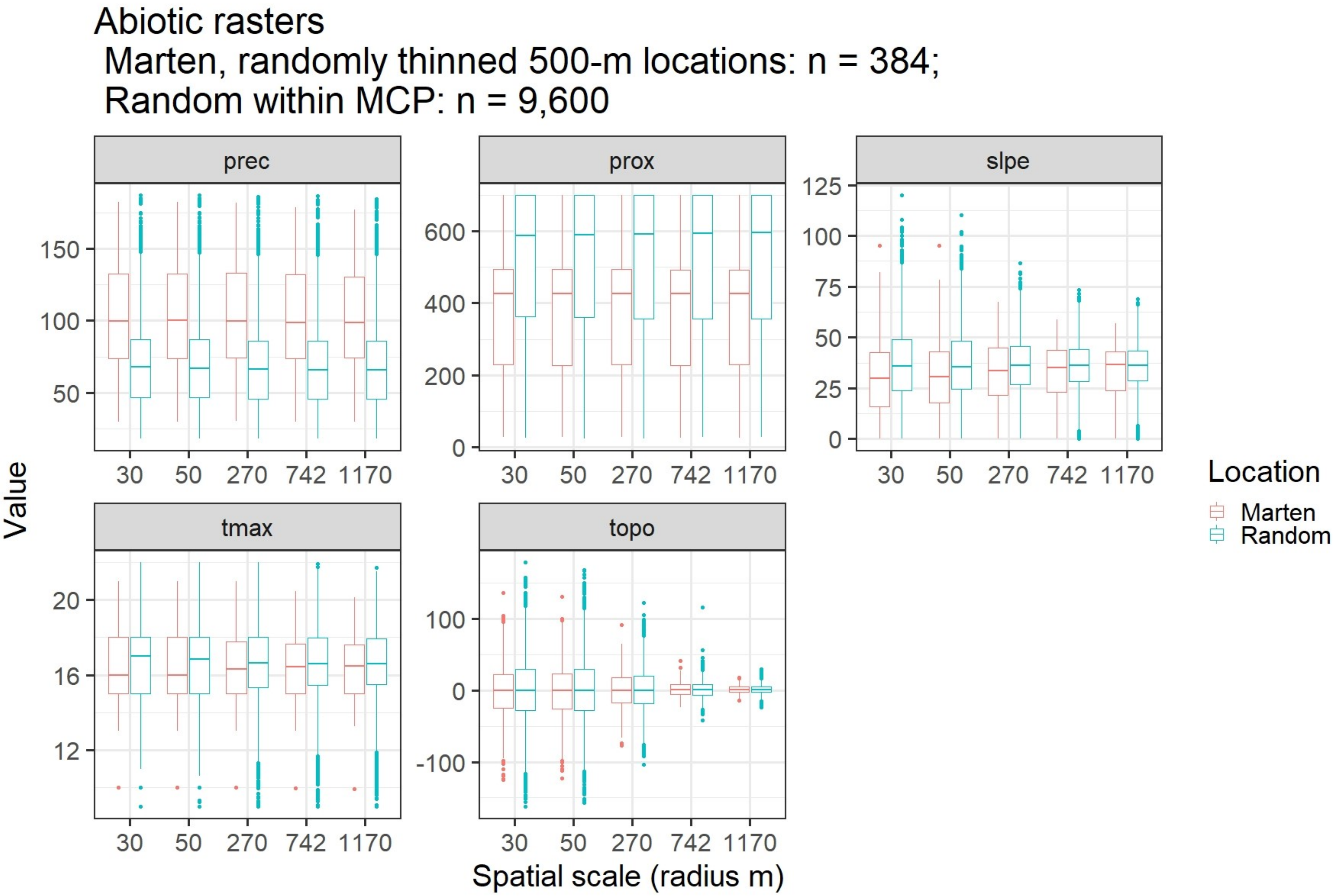

### Supplemental Figure S4

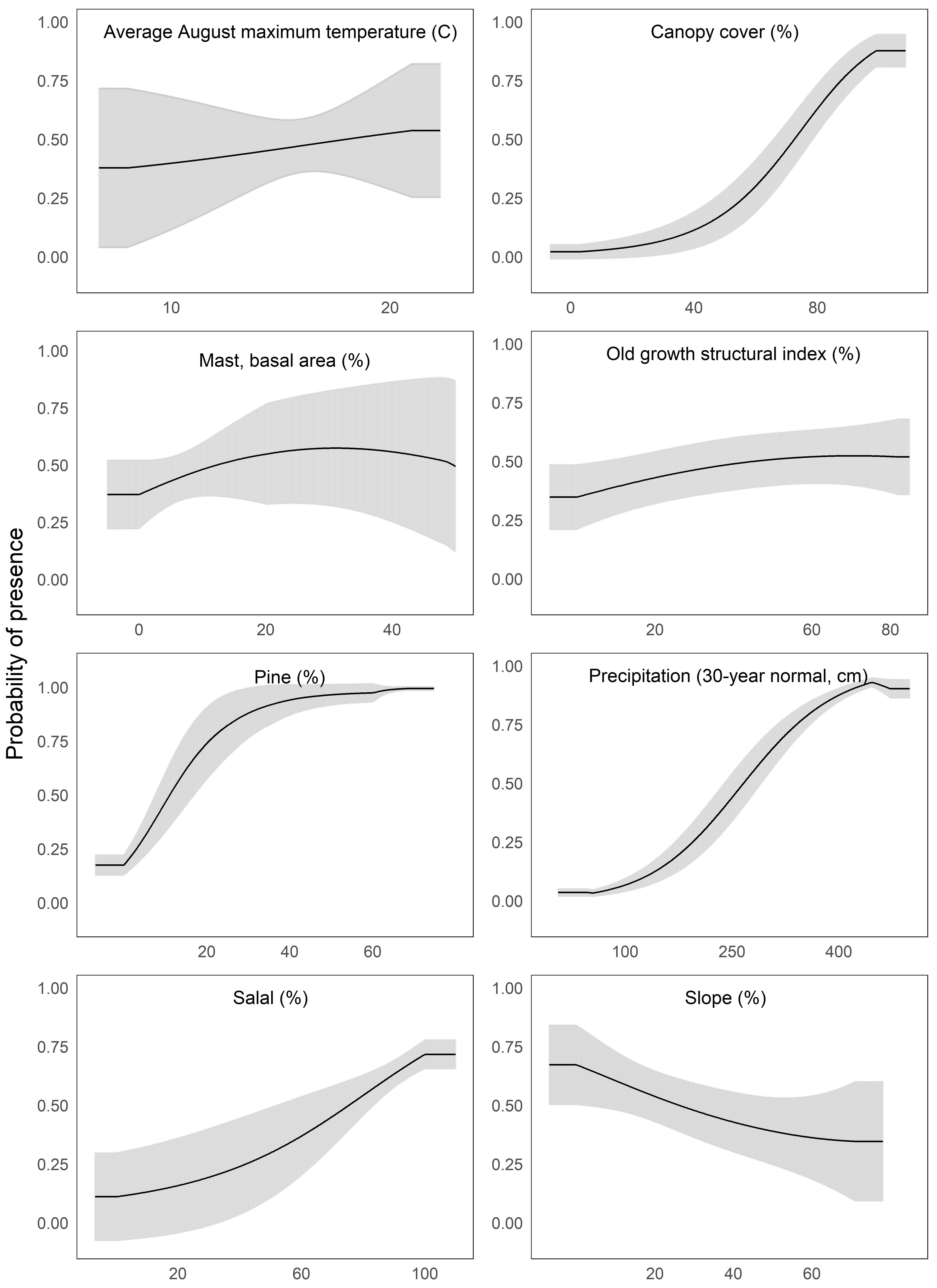
